# Long-read sequencing quantifies synthetic mRNA abundance, integrity and host response in vivo

**DOI:** 10.64898/2026.08.10.741180

**Authors:** Victoria M. McLeod, Daniel Yuen, Moore Z. Chen, Simone A. Beckham, Orlagh M. Feeney, Bruna Rossi Herling, Yifan Yang, Lara M. Molle, Pharvinderjit Kaur, Thomas J. Payne, Stewart A. Fabb, Colin W. Pouton, Christopher J.H. Porter, Angus P.R. Johnston

## Abstract

Synthetic mRNA can be used to reprogram biological systems and is increasingly used in vaccines, gene therapies and other advanced therapeutics. However, measuring mRNA abundance, molecular integrity and biological effects in complex samples remains challenging. Existing assays typically quantify short transcript regions or infer delivery from lipid or protein readouts. Here we present a long-read nanopore sequencing method that directly quantifies synthetic mRNA in complex cell and tissue samples. The approach enables absolute quantification of full-length synthetic mRNA, maps degradation at nucleotide resolution and simultaneously profiles associated host transcriptional responses. Applied to lipid nanoparticle (LNP) delivered mRNA in mice, the method revealed tissue-specific delivery and degradation patterns and uncovered a critical disconnect between mRNA accumulation and protein expression across organs. This approach enables integrated measurement of mRNA fate, integrity and biological responses, and will enable mechanistic studies of RNA delivery, stability, translation and innate immune recognition.

## Main

mRNA–lipid nanoparticle (mRNA/LNP) therapeutics are transforming disease prevention and treatment^1–4^. Beyond their success as vaccines, mRNA/LNPs are emerging as a platform for *in vivo* cell engineering and genome editing^5–7^. As these applications expand, understanding the delivery, biodistribution and reactogenicity to mRNA/LNP formulations is essential^8–10^. However, tracking them *in vivo* presents challenges not encountered with small-molecule drugs or protein therapeutics^11–13^. mRNA/LNPs are dynamic multicomponent systems where lipids and mRNA can separate after administration, translated protein provides an indirect and delayed measure of delivery, and single breaks render the RNA non-functional^14,15^.

Current biodistribution assays provide an incomplete picture of LNP/mRNA delivery. Quantification of lipid components by LC-MS/MS, radiolabelling or NMR/MRI^16,17^, does not capture the fate of the mRNA payload^18–21^. Protein-based readouts verify translation, but do not directly quantify mRNA delivery, as delivered transcripts may be retained, degraded or otherwise rendered poorly translated within specific cells or tissues. Probe-based assays such as RT–qPCR, digital PCR and branched-DNA hybridization are sensitive, but interrogate short transcript regions and therefore do not report RNA integrity^22,23^. Sequencing approaches have mainly been used to assess IVT mRNA quality before formulation rather than to quantify synthetic mRNA fate *in vivo*^24,25^.

To address these limitations, we developed a long-read sequencing method that directly quantifies synthetic mRNA in biological samples, distinguishes intact from degraded molecules and profiles host transcriptional responses from the same data. We benchmark the assay against electrophoresis and RT– qPCR, define sensitivity and multiplexing constraints, and apply it to LNP-delivered mRNA in mice. The resulting data reveal tissue-specific mRNA disposition and processing, and show that mRNA exposure and protein expression are not directly correlated.

## Results

### Nanopore sequencing improves resolution of mRNA integrity

We first tested whether nanopore cDNA sequencing (Fig 1a.) could measure IVT mRNA integrity with single-nucleotide resolution. We analysed Cre recombinase mRNA and compared the results with high-resolution microfluidic electrophoresis^26^. Direct RNA sequencing was not used because N^1^-methylpseudouridine interferes with current base-calling, and cDNA sequencing is faster and readily multiplexed. To accurately evaluate mRNA length, we developed an algorithm that excludes artificially truncated reads caused by premature sequencing termination or incomplete reverse transcription. This filtering is based on detection of the strand-switching primer, which is absent from incomplete cDNAs and prematurely terminated reads.

**Fig. 1:**
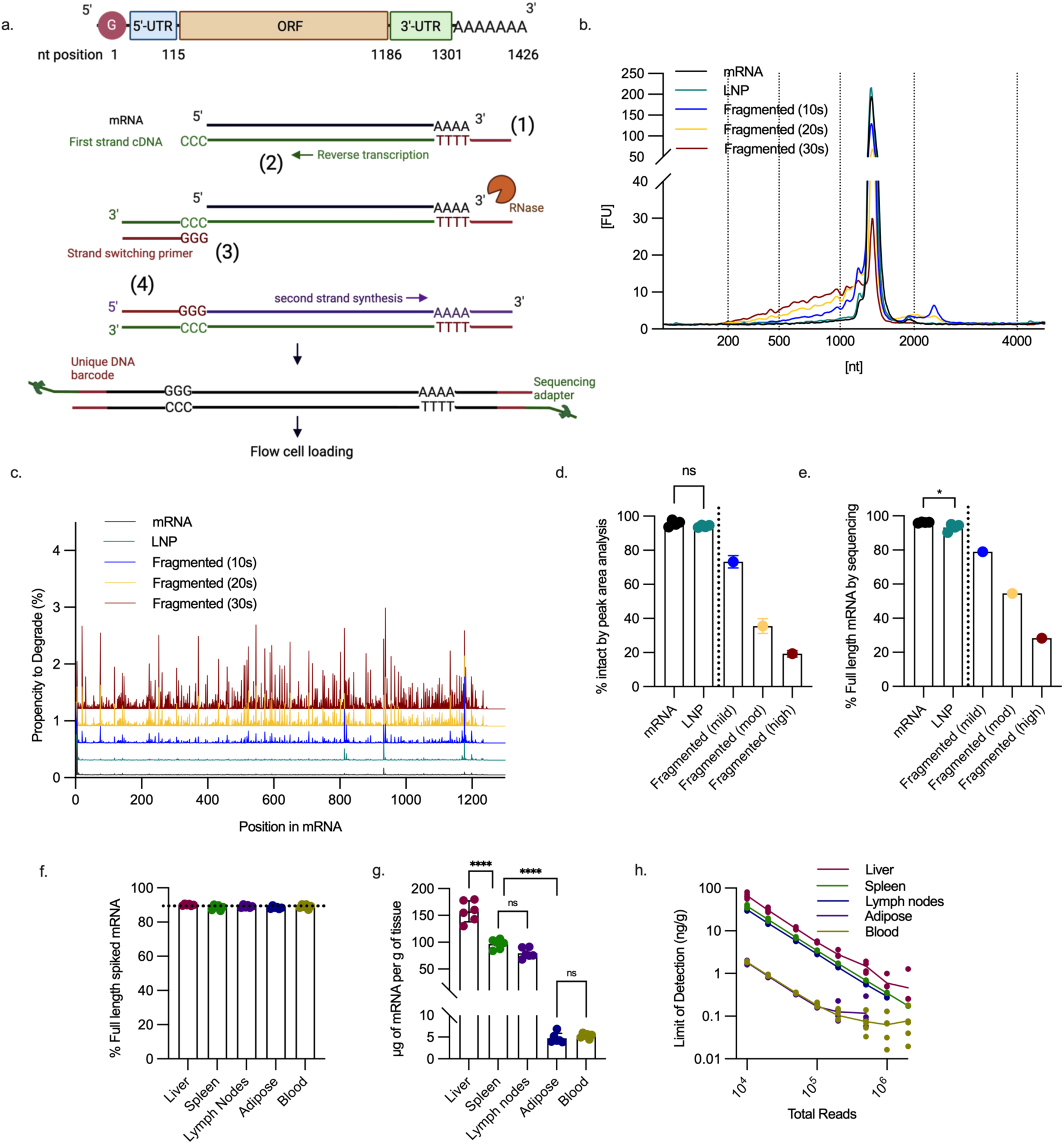
Nanopore sequencing accurately and quantitatively determines mRNA integrity in tissue. a, Schematic of full length Cre recombinase mRNA and nanopore cDNA sequencing library preparation. b, Bioanalyzer electropherograms of Cre mRNA before and after LNP encapsulation or simulated fragmentation. c, Positional fragmentation profile across the Cre mRNA sequence, showing the percent of fragmentation at each nucleotide location. Successive plots are vertically offset by 0.3% increments to prevent overlap and aid with visualisation. d, Primary peak area as a percentage of total electropherogram area and e, full length reads as a percentage of total reads. Data are mean ± s.d., n=4 independent LNP formulations and n=1 for fragmentated mRNA. *P<0.05 by two-way paired t-test. f, Full length reads as a percentage of total reads for LNP encapsulated mRNA spiked into tissues and processed through RNA extraction protocol. Horizontal dotted line indicates mRNA before tissue processing. Data are mean ± s.d., tissue from n=6 mice. g, Endogenous cellular mRNA content of different organs. Data are mean ± s.d., tissue from n=6 mice. ****P<0.0001 by repeated measures one-way ANOVA with Tukey’s multiple comparison between tissues. h, Limit of detection (LOD) estimated from false positive reads in PBS dosed mouse tissues across increasing sequencing depth (number of reads). LOD was defined as the mean + 3 s.d. false positive reads from tissue of n=25 mice.

Microfluidic electrophoreses (Bioanalyzer) showed a dominant Cre mRNA peak at ∼1426 nt, corresponding to 97±2% of the mRNA, while nanopore sequencing identified 96 ± 0.3% full-length (spanning the 5′ cap to the start of the poly(A) tail) reads (Fig. 1b-e). To confirm nanopore sequencing could consistently detect degraded mRNA, we used Mg^2+^-mediated RNA hydrolysis to generate increasingly fragmented species of Cre mRNA (following 10, 20, 30s reaction times). Microfluidic electrophoresis showed a progressive decrease in the full-length peak and accumulation of shorter products (Fig. 1b). However, overlap between the full-length peak and a shoulder of shorter fragments limited accurate quantification. By estimating the area of the peak at ∼1426 nt, full-length mRNA dropped to 19% after 30 seconds of fragmentation (Fig. 1d). Nanopore sequencing showed a similar loss of full-length mRNA, with 28% full length remaining after 30 seconds (Fig. 1e).

Nanopore sequencing also showed that fragmentation was non-uniform across the transcript (Fig. 1c), a feature not evident from the corresponding electropherogram. Hydrolysis for 10, 20 and 30 s produced similar fragmentation patterns, with fragmented products increasing in abundance relative to the full-length transcript. We also assessed RT–qPCR as a measure of mRNA integrity using primer sets spanning 97.7% of the Cre mRNA sequence (Supplementary Fig. 1a–h). However, RT–qPCR incorrectly detected an eightfold higher apparent abundance of full-length mRNA in the 30-s fragmented sample than in the unfragmented control (Supplementary Fig. 1f). This likely reflects off-target amplification from shorter RNA fragments, which is consistent with the reduced melting temperature of amplicons generated from the fragmented mRNA compared with full-length mRNA (Supplementary Fig. 1h).

We next examined whether LNP encapsulation affected mRNA integrity using the Moderna Spikevax LNP formulation (Fig. 1c,e; Supplementary Fig. 2a-h). Nanopore sequencing detected a modest but statistically significant reduction in full-length mRNA after encapsulation (93 ± 2.1%; P = 0.0489; Fig. 1e). In contrast, microfluidic electrophoresis did not resolve a statistically significant decrease (94 ± 0.7%; P = 0.1630; Fig. 1d). The degradation profile of LNP-encapsulated mRNA was not random, with discrete cleavage hotspots across the transcript (Fig. 1c). One hotspot at nucleotide position 938 was present in both unencapsulated and LNP encapsulated mRNA, but was more pronounced after encapsulation. Additional hotspots at positions 820, 825 and 1177 showed limited cleavage in unencapsulated mRNA, but exceeded 0.1% cleavage frequency after LNP encapsulation. Together, these data show that nanopore long-read sequencing can quantify synthetic mRNA integrity and identify sequence positions that are particularly susceptible to degradation.

### Nanopore sequencing can accurately quantify synthetic mRNA in tissue by referencing endogenous mRNA

We next validated our sequencing method in complex tissue samples. To determine whether sample processing altered mRNA integrity, we spiked mouse tissues with known amounts of LNP-encapsulated mRNA or deliberately fragmented mRNA before tissue homogenisation, RNA extraction and library preparation. Tissue processing did not measurably reduce mRNA integrity (Fig. 1f, Supplementary figure 2i), and degradation profiles from fragmented spike-ins were preserved (Supplementary figure 2j). This indicates the workflow recovers both intact and degraded synthetic mRNA species from complex biological matrices.

The number of synthetic mRNA reads cannot be used to infer absolute abundance, as the number of reads is dependent on sequencing depth. The ratio of synthetic to endogenous mRNA reads provides a measure of relative abundance, but not absolute quantitation. To enable quantification, we spiked a known amount of synthetic mRNA into each tissue and used the resulting synthetic-to-endogenous read ratio to estimate the endogenous mRNA content per g of tissue (Fig. 1g, Supplementary Table 1). Once established for each tissue, this reference value can be used to quantify synthetic mRNA abundance in unknown samples.

### Sensitivity and limit of detection of synthetic mRNA quantitation in tissue

Nanopore sequencing sensitivity is determined by sequencing depth. To balance sensitivity against sequencing cost and time, we assessed how the limit of detection (LOD) for synthetic mRNA varied with increasing sequencing depth. We sequenced RNA from liver, spleen, lymph nodes, adipose tissue, and blood from PBS-treated control mice to estimate tissue specific false-positive rates. For each tissue, the LOD was defined as the mean false-positive counts plus three standard deviations (n = 25 mice), with a minimum threshold of three reads. For liver, spleen, and lymph nodes, 50,000 total reads were sufficient to detect synthetic mRNA at concentrations ≥10 ng g^-1^ of tissue (Fig. 1h). In blood and adipose tissue, which contain lower amounts of endogenous mRNA, 20,000 reads were sufficient to detect synthetic mRNA at ≥1 ng g^-1^ tissue. Increasing sequencing depth to 500,000 reads improved sensitivity in liver, spleen, and lymph nodes, enabling detection of synthetic mRNA at levels >1 ng g^-1^.

Using RT-qPCR to probe a near full-length Cre mRNA amplicon showed reduced sensitivity of 2 and 20 ng g^-1^ in liver and spleen, respectively (Supplementary Table 2). By comparison, RT-qPCR using an optimised 104 base amplicon quantified Cre mRNA in liver and spleen down to 0.02 and 0.05 ng g^-1^, respectively (Supplementary Table 2). Thus, although short-amplicon RT–qPCR provides higher analytical sensitivity, its performance depends strongly on primer design and amplicon length, whereas nanopore sequencing directly measures transcript length and integrity.

### Nanopore sequencing reveals tissue-specific biodistribution and kinetics of intact mRNA

Having confirmed nanopore sequencing accurately quantifies total synthetic mRNA, percent of full-length synthetic mRNA and degradation profiles in complex tissues, we applied this method to track mRNA biodistribution in mice over 24 h. Ai14 tdTomato reporter mice received 500 µg kg^-1^ Cre mRNA/LNPs via intravenous injection. Blood was collected before perfusion, and liver, spleen, lymph nodes and adipose tissue were analysed for mRNA abundance, integrity and degradation profiles (Fig 2a).

**Fig. 2:**
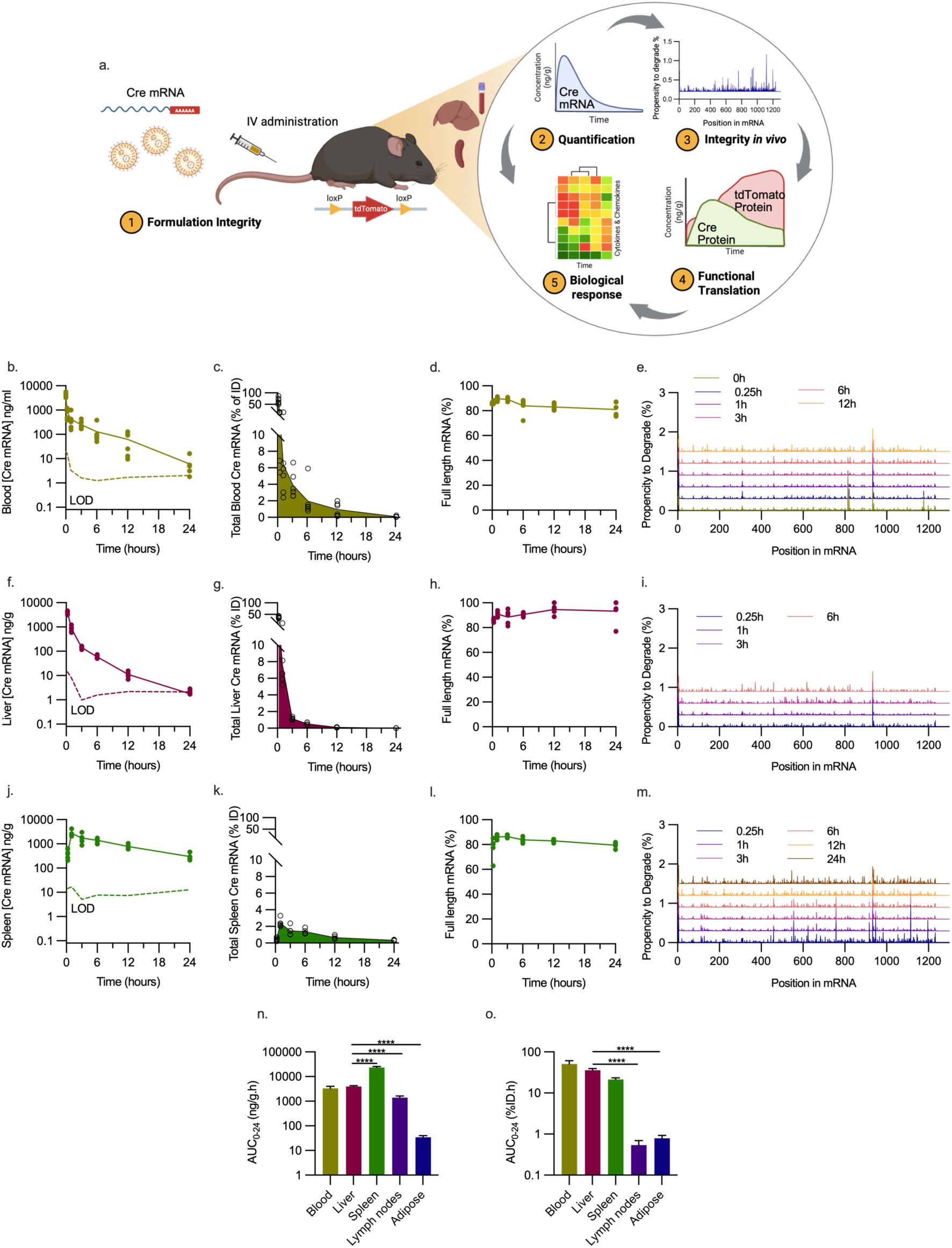
Tissue specific pharmacokinetics reveal nonuniform distribution, elimination and processing of LNP delivered mRNA. **a**, Study design and analytical outputs in Ai14 tdTomato reporter mice dosed with Cre mRNA. **b-m**, Nanopore quantification of Cre mRNA in blood (b-e), liver (f-i), and spleen (j-m), shown as mass per volume with the limit of detection (LOD) determined from PBS-dosed mice (b,f,j) or percentage of injected dose (%ID) in total organ (c,g,k), mRNA integrity expressed as percent of sequences that are full length (d,h,i), and propensity for degradation at each nucleotide position in Cre mRNA(e,i,m). Degradation analysis at 24 h for blood, 12 and 24 h for liver were excluded due to insufficient reads (n=6 mice at 0.25h and n=5 all other time points). **n**, Area Under Curve (AUC) calculated from mRNA concentration-time plots ± SE. \**p*<0.0001 compared to liver following Welch one-way ANOVA. **o**, Area Under Curve (AUC) calculated from %ID over time plots ± SE. \**p*<0.0001 compared to liver following Welch one-way ANOVA.

Cre mRNA was detected at high levels in the blood (4,312 ± 898 ng mL^-1^), representing 66 ± 14% of the injected dose (%ID) 30s after injection (Fig. 2b,c). Circulating mRNA followed classic two-phase pharmacokinetic behaviour, with rapid distribution from blood (T_1/2_ 4.6 min, 95% CI 0-11 min), followed by slower terminal elimination (T_1/2_ 235 min, 95% CI 197-292 min; Supplementary Table 3). By 24 h, <1% of the injected dose remained in circulation. Interestingly, the full-length fraction (81 ± 5%) and degradation profile were consistent over the 24 h sampling period (Fig. 2d,e). Because plasma is RNase rich^27^, this stability is consistent with circulating mRNA remaining encapsulated in LNPs and protected from degradation.

Standard LNP formulations are effectively taken up by the liver through interactions with serum lipoproteins and their receptors present on hepatocytes^28^. As expected, Cre mRNA accumulated rapidly in the liver, quantified at 4,050 ± 489 ng g^-1^ (38 ± 4.0 %ID) after 15min (Fig. 2f,g). mRNA concentration declined in a biphasic manner with an initial rapid phase (T_1/2_ of 17 min; 95% CI 14-21 min) and a second slower phase (T_1/2_ 142 min; 95% CI 77-317 min). After 24 hours, the concentration of mRNA present was at the limit of detection (1.8 ± 1.1 ng g^-1^; Fig 2b). The persistently high full-length fraction (>90%) suggests that hepatic mRNA degradation and clearance occur together rather than through accumulation of stable degradation intermediates (Fig 2h,i)^29,30^.

The spleen, another major reticuloendothelial organ, showed notably different mRNA kinetics, with slower accumulation and longer retention (Fig 2.j,k). Cre mRNA levels peaked at 1h (2,707 ± 866 ng g^-1^, 2.4 ± 0.5 %ID), then slowly declined with a single elimination phase (T_1/2_ 303 min; 95% CI 171-615 min; Fig 2j). At 24 h, splenic Cre mRNA (300 ± 93 ng g^-1^, 0.3 ± 0.1 %ID ) was ∼100 fold higher than in the liver. Although the full-length fraction in the spleen was similar to other tissues (80-90%; Fig. 2l), the degradation profile showed a different pattern (Fig 2m). Notably, we observed consistently increased degradation at several positions within the Cre mRNA (555, 665, 758, 917, and 1115 nt positions from 5’ end; Fig. 2m). These data indicate tissue-specific mRNA processing in spleen and liver.

Lymph nodes, which are compositionally similar to the spleen, contained less Cre mRNA per gram of tissue (Supplementary Fig. 3a). At 1 h, the lymph nodes contained ∼5% of the Cre mRNA observed in the spleen, but showed similar retention kinetics and full-length representation over time (Supplementary Fig. 3b,c). Adipose tissue had the lowest Cre mRNA content per tissue mass (Supplementary Fig. 3d-f).

To quantify organ exposure, we calculated the 24 h area under the curve (AUC, Fig. 2n,o). On a mass-normalized basis, mRNA exposure was ∼ six-fold higher in the spleen compared to the equivalent mass of liver (*p*<0.0001, Fig. 2n). This highlights the distinct differences in how these two major organs process LNPs and mRNA cargos. The AUC for %ID was more similar in the spleen compared to the liver (21 vs. 36 %ID.h, respectively, *p*=0.0545, Fig. 2o), reflecting the ∼10-fold greater mass of the liver compared to spleen. Nanopore measurements aligned with RT–qPCR across liver and spleen time courses, while providing the additional integrity and degradation information unavailable from RT-qPCR (Supplementary Figure 3g,h and Supplementary Table 4).

### mRNA biodistribution does not correlate with protein expression

We next examined how well mRNA exposure predicts functional protein output. Reporter protein expression is widely used as a proxy for assessing LNP-mediated mRNA delivery^31^. However, it reports successful translation, and does not determine how much mRNA cargo is delivered to tissues nor how efficiently cellular uptake or translation occurs. In Ai14 reporter mice, translation of Cre recombinase excises a floxed STOP cassette, permanently activating tdTomato transcription^32^. We therefore interrogated three linked outputs of this system across tissues: (i) Cre protein abundance by ELISA, (ii) Cre-mediated tdTomato transcript induction by nanopore sequencing, and (iii) tdTomato reporter expression by ELISA and fluorescence. This allowed comparison of mRNA exposure with both immediate protein production and downstream reporter activation.

Cre protein was undetectable in plasma (LOQ = 0.17ng ml^-1^). In liver, Cre protein was detectable within 15 min and peaked 3-6 h after administration (5,200 ± 1,000ng g^-1^ at 3h, Fig. 3a). tdTomato mRNA and protein accumulated over 24 h (Fig. 3b,c). Liver sections collected 6 h after dosing showed extensive tdTomato expression in hepatocytes, whereas Kupffer cells showed low or undetectable expression (Fig. 3d). By 24 h, strong hepatocyte fluorescence masked signals from immune cells, illustrating the difficulty of quantifying cell-specific expression using fluorescent reporters. Flow cytometry analysis of resident liver immune cells detected tdTomato predominantly in phagocytic cells, including monocytes (53.1 ± 5.4% positive) and Kupffer cells (5.2 ± 2.3% positive), which represented ∼11% and 2% of total liver CD45^+^ immune cells, respectively (Fig 3e, Supplementary Figure 4). Hepatocytes were not quantified by flow cytometry because of their fragility.

**Fig. 3:**
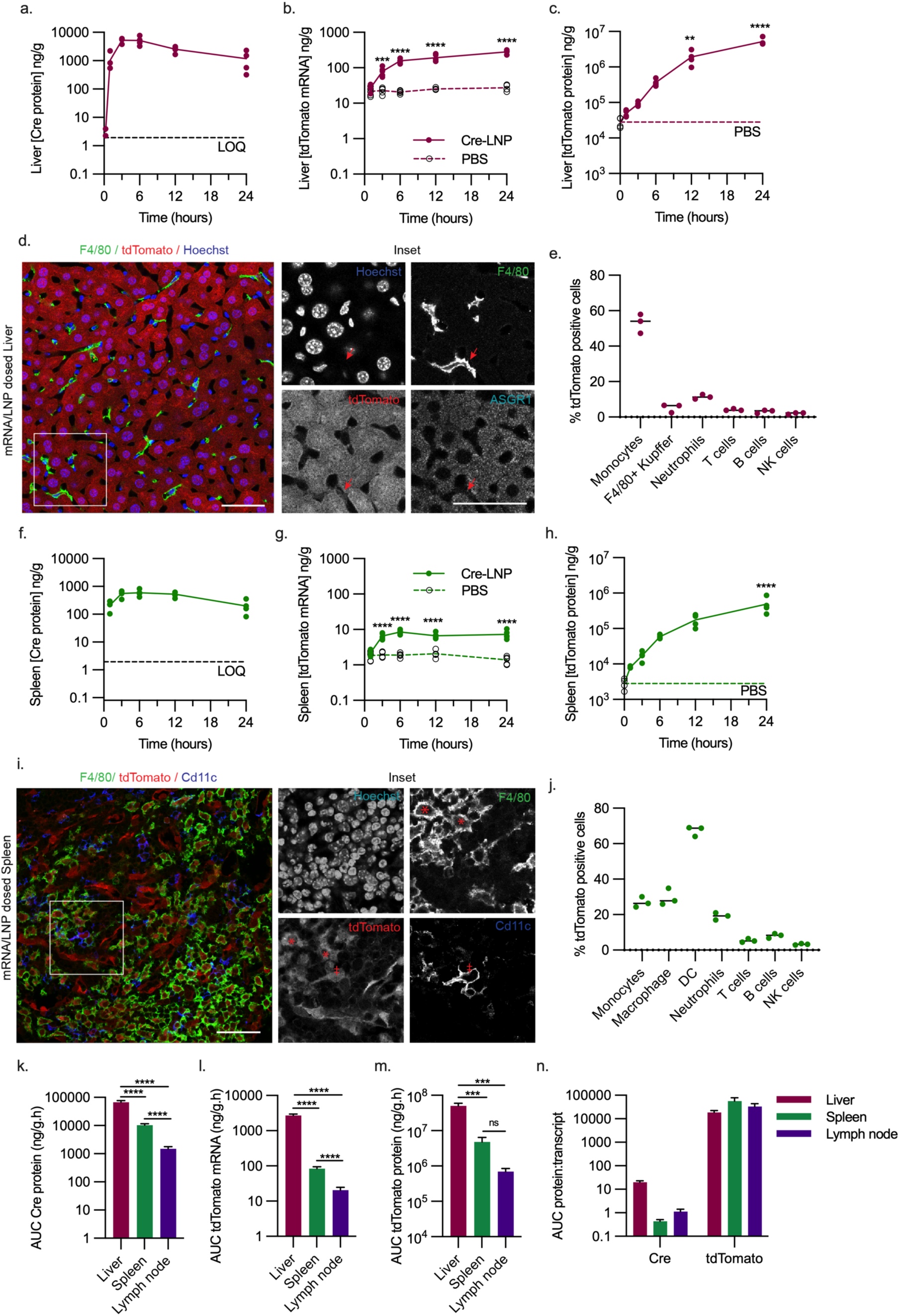
Protein level readouts do not correlate with mRNA delivery. a–j, Protein expression and reporter activation in liver (a–e) and spleen (f–j) after Cre mRNA/LNP administration in Ai14 tdTomato reporter mice. Cre protein quantified by ELISA in liver (a) and spleen (f), with the limit of quantification (LOQ) determined from spiked tissue standards shown as a dashed line; n = 4 mice per time point. tdTomato reporter transcript abundance quantified in liver (b) and spleen (g) after Cre mRNA/LNP (filled symbols, solid line) or PBS administration (open symbols, dashed line); n = 5 mice per time point; ***P < 0.001, ****P < 0.0001 by two-way ANOVA with Sidak’s multiple-comparison test. tdTomato protein quantified by ELISA in liver (c) and spleen (h), with PBS controls defining baseline expression at t = 0 (open symbol; dashed line, mean); n = 4 mice per time point; **P < 0.01, ****P < 0.0001 by one-way ANOVA with Dunnett’s multiple-comparison test versus t = 0. Liver (d) and spleen (i) sections collected 6 h after Cre mRNA/LNP administration show tdTomato fluorescence with staining for nuclei (Hoechst), macrophages (F4/80) and hepatocytes (ASGR1; liver) or dendritic cells (CD11c; spleen). Insets show individual channels; arrow indicates an F4/80-positive macrophage in liver, asterisk indicates an F4/80-positive macrophage in spleen and double dagger indicates a tdTomato-positive CD11c-positive dendritic cell. Scale bars, 50 µm. tdTomato-positive immune cells were quantified by flow cytometry in liver (e) and spleen (j) 24 h after Cre mRNA/LNP administration; n = 3 mice. k–m, Area under the curve (AUC) over 24 h for Cre protein (k), tdTomato transcript abundance (l) and tdTomato protein expression (m) across tissues. Data are mean ± s.e.; ***P < 0.001, ****P < 0.0001 by Welch one-way ANOVA with Dunnett’s T3 multiple-comparison test. n, Protein-to-mRNA transcript ratios for Cre and tdTomato across tissues, calculated from AUC values with propagated absolute error (s.e.).

The spleen produced less Cre protein (595 ± 169 ng g^-1^ at 6h; Fig. 3f) than the liver, despite sustained mRNA exposure. tdTomato transcripts plateaued by 6 h (Fig. 3g), whereas tdTomato protein continued to accumulate (Fig. 3h). This suggests the population of cells that are exposed to Cre undergo recombination within the first 6 h. The continued presence of Cre protein within the spleen tissue does not seem to activate additional cells. In contrast, liver tdTomato transcripts progressively increased, consistent with activation of additional cells over time. Spleen sections collected 6 h after dosing showed sparser tdTomato expression than liver (Fig. 3i). tdTomato signal was concentrated in red pulp zones, including in both macrophages and dendritic cells (Fig. 3i), while it was largely absent from white pulp zones (Supplementary Figure 5). Flow cytometry confirmed high expression in phagocytic cells (Fig. 3j), including dendritic cells (67 ± 3%), macrophages (30 ± 5%) and monocytes (27 ± 3%) although each only accounted for ∼0.5-1.5% of CD45^+^ cells.

Lymph node Cre protein peaked 3-12h after administration but at ∼5-fold lower levels than spleen, and was undetectable in adipose tissue (Supplementary Figure 6a). Both lymph node and adipose showed detectable tdTomato protein by 24h, despite adipose having no detectable transcript elevation (Supplementary Figure 6b-f).

Across tissues, protein output did not scale with mRNA exposure. Liver Cre protein exposure, calculated as the AUC of the concentration-time profile, was 6.5-fold higher than the spleen (67,698 ± 9,699 vs 10,274 ± 1,239 ng g^-1^ h respectively, Fig. 3k), despite 5.9-fold lower mRNA exposure (Fig 2t). This disconnect shows that reporter protein alone is insufficient for modelling mRNA pharmacokinetics or functional delivery. The liver also showed higher tdTomato reporter transcript (Figure 3l) and protein expression (Figure 3m) compared to spleen, consistent with Cre protein levels (but not mRNA). Blood and adipose tissue were excluded because Cre protein was below the limit of detection. This reinforces the conclusion that synthetic mRNA is processed differently across tissues. Direct comparisons of protein-to-mRNA ratios between genes remain constrained by protein stability and turnover; nevertheless, the protein-to-mRNA ratio for endogenously expressed tdTomato was 10S– 10‘-fold higher than that for Cre expressed from synthetic mRNA, indicating substantially lower output from delivered transcripts (Fig. 3n). Together, these findings show that indirect readouts, including protein expression alone, are insufficient to model mRNA pharmacokinetics or functional delivery.

### Nanopore transcriptomics identifies host response pathways activated by mRNA/LNP delivery

LNP formulations are known to stimulate the immune system, which can be advantageous for vaccines, but detrimental for other therapeutic applications^33^. By capturing synthetic mRNA and endogenous transcripts in parallel, long-read sequencing enables simultaneous analysis of mRNA fate and formulation reactogenicity. We compared cytokine and chemokine transcripts measured by nanopore sequencing with protein abundance measured by multiplex bead assay. In spleen, transcripts and proteins associated with acute inflammation and innate immunity increased after LNP/mRNA administration (Fig. 4a). Transcript-level responses generally peaked before protein responses and were consistently higher than matched tissue protein changes. Higher transcript changes are possibly because secreted cytokines redistribute into plasma. Pearson’s correlation analysis of transcript and protein fold changes were strongly correlated across time points, particularly at 12 h (Fig. 4b, Supplementary Figure 7). Plasma protein profiling showed broad cytokine induction, whereas whole-blood cellular transcripts were limited mainly to *Cxcl10* and *Il1b* (Fig 4c). Liver, adipose and lymph nodes, showed similar upregulation of cytokine and chemokine transcripts to the spleen. Tissue transcriptomics therefore localized the response and were not restricted to a predefined analyte panel (Fig. 4c,d).

**Fig. 4:**
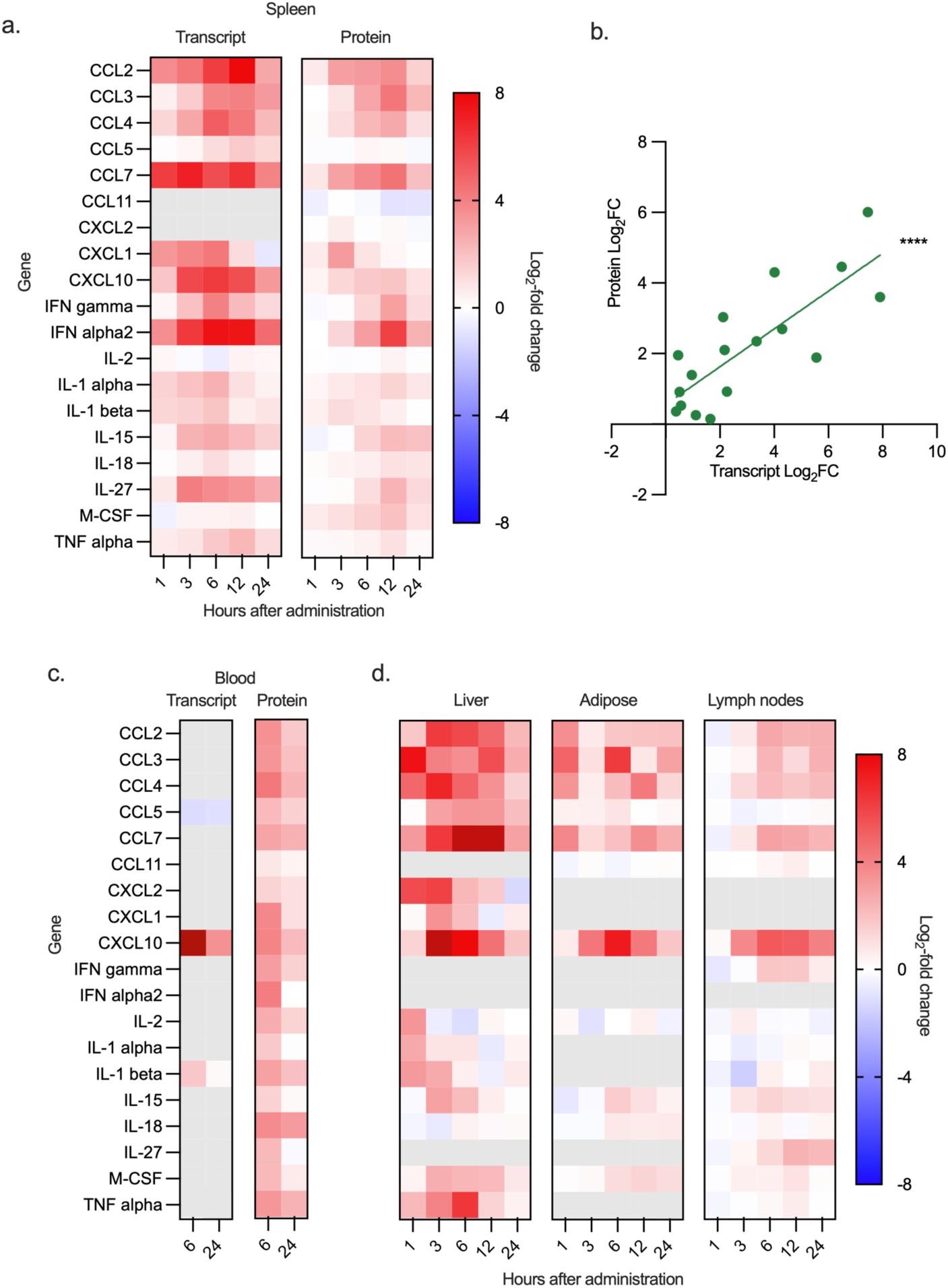
Nanopore-based cytokine profiling correlates to protein-level inflammatory responses. a, Heatmap showing log2-fold changes in transcript abundance measured by nanopore sequencing and protein abundance measured by multiplex bead assay for 19 cytokines and chemokines in spleen. Targets include mediators of innate inflammation, adaptive immune regulation and antiviral responses. Fold changes were calculated at each post-administration time point relative to PBS-dosed controls. b, Correlation between cytokine transcript and protein log2-fold changes in spleen 12 h. ****P < 0.0001 by Pearson correlation; r = 0.8058. c, Comparison of cytokine transcript abundance in whole blood and protein abundance in plasma at 6 and 24 h after mRNA/LNP administration. d, Heatmap showing log2-fold changes in transcript abundance in liver, adipose tissue and lymph nodes. For heatmaps, red indicates upregulation, blue downregulation, white no change relative to PBS-dosed controls and grey undetected transcripts. n = 5 mice for transcript and protein analysis.

The large RNA-seq data sets can provide a breadth of information to probe complex biological pathways in response to the delivery of mRNA. Differential expression and KEGG enrichment extended this analysis beyond cytokines. Multiple immune pathways, including antigen processing, cytokine signalling, Toll-like receptor signalling and RIG-I-like receptor signalling, were enriched across tissues after LNP/mRNA delivery (Fig. 5). Liver and blood also showed sustained downregulation of ribosomal transcripts, consistent with interferon-associated translational shutoff,^34^ whereas spleen did not, reinforcing organ-specific differences in response to foreign mRNA and LNPs.

**Fig. 5:**
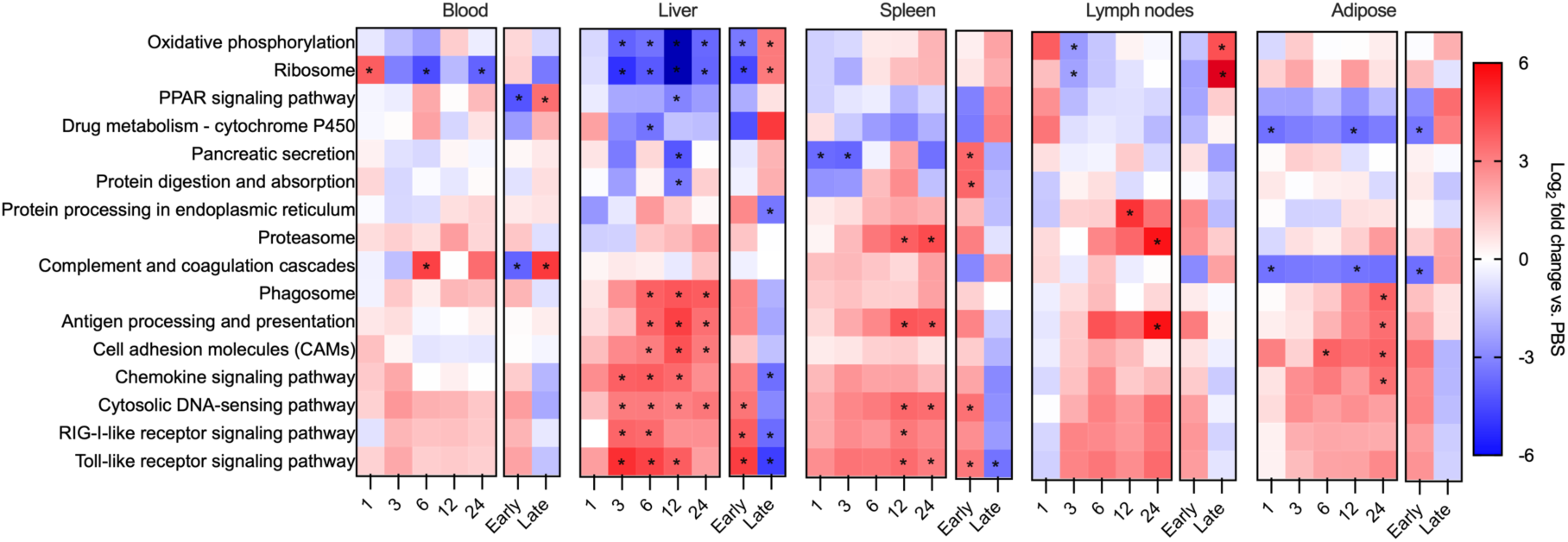
KEGG pathway analysis identifies transcriptional responses to LNP/mRNA Delivery. Differentially abundant transcripts detected by nanopore sequencing were assigned to KEGG pathways, and pathway-level fold changes were calculated relative to PBS-dosed tissues. Heatmaps show pathway fold changes at each sampled time point after mRNA/LNP administration, together with trend analyses summarizing the initial response over the first 6 hours (early) and from 6-24 hours (late). *FDR < 0.05.

Finally, we examined genes encoding RNA-sensing and antiviral effector proteins. Endosomal and cytosolic RNA sensors, including *Tlr3* (TLR3), *Ifih1*(MDA5), *Pkr* (PKR), *Zbp1* (ZBP1) and *Rigi* (RIG-I), were induced in multiple tissues (Fig. 6 and Supplementary Figures 8-11). The OAS/RNase L axis and *Zc3hav1* (ZAP) were also upregulated, whereas *Dicer1* (DICER1) changed little, suggesting that classical antiviral RNA-degradation pathways rather than RNA interference dominate the response^35^. Translation-inhibitory *Ifit* genes were induced broadly, consistent with cellular restriction of exogenous RNA translation. Together, these analyses show that our nanopore-sequencing method can connect synthetic mRNA abundance and integrity with tissue-level innate immune mechanisms.

**Fig. 6:**
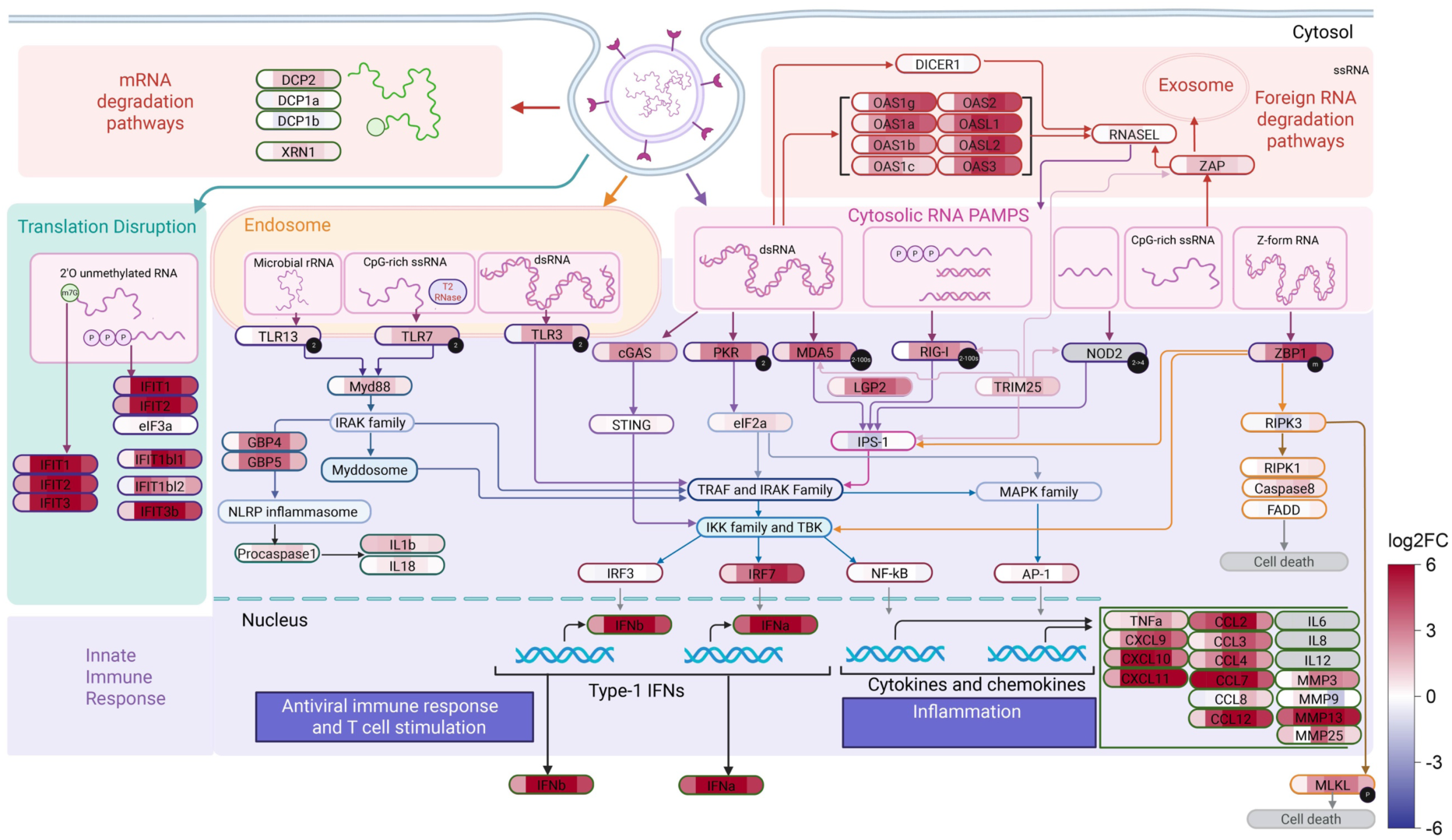
mRNA/LNP delivery induces RNA-sensing, interferon and antiviral effector pathways. Schematic representation of RNA-sensing and effector pathways upregulated in spleen after mRNA/LNP administration. Heatmap overlays indicate gene-level upregulation, shown as log2-fold change in transcript abundance relative to PBS-dosed controls.

## Discussion

We have developed a long-read nanopore-sequencing method that quantifies synthetic mRNA biodistribution, molecule-level integrity and host transcriptional responses from the same biological samples. This approach provides an integrated method to probe the key parameters of therapeutic mRNA pharmacology.

Compared with microfluidic electrophoresis, long-read sequencing provides a more accurate and precise estimation of full-length mRNA and uniquely identifies the nucleotide positions most susceptible to degradation. Microfluidic electrophoresis is unable to resolve degradation sites and is incompatible with complex biological matrices, whereas long-read sequencing retains high resolution across all tissues tested. While RT-qPCR is the current gold standard approach for quantification of mRNA, it is reliant on single target primer or probe design, and is only reliable for amplifying short-amplicons. We showed good alignment of mRNA quantitation in tissue with long-read sequencing when compared to RT-qPCR with optimised primers. Although less sensitive, long-read sequencing yields substantially richer information, including full degradation profiles, quantification of intact transcripts, and comprehensive transcriptome-wide measurements which cannot be achieved with RT-qPCR.

Applied *in vivo*, our method revealed organ-specific mRNA behaviour. Liver showed rapid mRNA accumulation and clearance with efficient protein production, whereas spleen retained mRNA for longer but translated it less efficiently. This disconnect indicates that mRNA exposure, molecular integrity and tissue translation capacity must be measured separately when evaluating delivery systems. Importantly it is apparent that protein expression is a poor indicator of mRNA delivery. As such interpretation of LNP formulations for mRNA delivery based on protein expression read outs is intrinsically flawed.

The endogenous transcriptome component provides an additional readout of formulation reactogenicity and biological behaviour. Cytokine transcript changes correlated strongly with protein-level inflammatory responses. Furthermore, pathway analysis of the transcriptional changes is an unbiased approach that identified RNA-sensing, antiviral and translational-shutoff signatures that differed across tissues. These data are useful both for screening delivery formulations and for mechanistic studies of innate recognition of therapeutic RNA.

Overall, long-read sequencing provides a multidimensional understanding of therapeutic mRNA delivery, which cannot be achieved with existing analytical methods. By integrating absolute quantification, positional integrity mapping, protein correlation, and transcriptome profiling, this approach offers an analytical platform for evaluating mRNA therapeutics *in vivo*. The ability to resolve both payload fate and host response at high resolution will be valuable for optimising LNP formulations, improving therapeutic design, and advancing next-generation mRNA medicines.

## Methods

### Messenger RNA and LNP synthesis

mRNA encoding Cre recombinase fused to a nuclear localisation sequence was synthesized as previously described^36^. Lipid nanoparticles were prepared by microfluidic mixing on a NanoAssemblr™ Ignite (Precision Nanosystems). A lipid mixture matching the composition of the Moderna COVID-19 vaccine containing SM-102 (DC Chemicals #DC52025), DSPC (Avanti Polar Lipids #850365), cholesterol (Merck #C8667), PEG2000-DMG (Avanti Polar Lipids #880151), at a molar ratio of 50:10:38.5:1.5 was dissolved in ethanol. In parallel, in vitro transcribed (IVT) mRNA was diluted in 0.3M sodium acetate buffer (pH 4). Lipids and mRNA were combined at 3:1 flow rate ratio (aqueous:organic) under continuous flow, yielding LNPs with an N/P ratio of 6. Formulations were dialyzed against PBS to remove ethanol, then sterile-filtered and stored at 4 °C until use. Consistency in particle size and encapsulation efficiency was confirmed by dynamic light scattering and RiboGreen (Thermo Fisher) assay, respectively.

### mRNA fragmentation and microfluidic electrophoresis

Simulation of mRNA fragmentation was conducted using NEBNext® Magnesium RNA Fragmentation Module (NEB #E6150S) according to manufacturer instructions. Synthetic mRNA was incubated in fragmentation buffer for 10, 20 and 30 seconds at 94°C in a preheated thermocycler before immediate addition of stop solution on ice. To release mRNA encapsulated in LNPs, 1-2ug mRNA/LNP was incubated in 1/3 volume of 2% triton-X 100 for 10 mins at 37°C. All reaction clean-up was performed using 1:1 volume of Mag-Bind® TotalPure NGS magnetic beads (Omega Bio-tek) and 70% ethanol wash before elution in nuclease-free water to achieve desired concentration. Microfluidic electrophoretic assessment was performed on an Agilent Bioanalyzer using RNA 6000 nano kit with 25-200ng of Cre mRNA loaded. Peak area and size were determined using 2100 Expert Software (Agilent Technologies) with manual peak integration.

### Animals & Biodistribution Studies

All experiments were conducted in accordance with the Australian Code for the Care and Use of Animals for Scientific Purposes and approved by the Monash Institute of Pharmaceutical Sciences Ethics committee (approvals 37404 and 37898). Transgenic Ai14 mice (B6.Cg-Gt(ROSA)26Sortm14(CAG-tdTomato)Hze/J, stock number 007914) were purchased from Jackson Laboratory (Bar Harbor, ME, USA) and maintained at Monash Animal Research Platform. C57BL/6J mice were supplied by Monash Animal Research Platform. For all experiments male and female mice were used, aged 6-14 weeks, maintained on a 12 h light/dark cycle with access to water and food at all times.

Ai14 mice were dosed intravenously with 0.5 mg/kg LNP encapsulated Cre mRNA and killed at 0.25, 1, 3, 6, 12 or 24h post-administration. An additional t=0 blood sample was obtained by submandibular bleed immediately following dose administration in the 0.25 h group. At termination blood was collected via cardiac puncture into EDTA K2 tubes and 0.2 ml whole blood was mixed with QIAzol® (Qiagen) and frozen. Mice were injected with 100mg/kg sodium pentobarbitone and cardiac perfused with PBS containing 10mM EDTA and organs rapidly snap frozen in liquid nitrogen. For biodistribution studies, data represents n=5-6 mice from 3 independent LNP formulation dosed cohorts. Within each dosing cohort, available mice were assigned into time points and dosing groups within cages in an attempt to balance sex and litter representation where possible. No predetermined randomisation strategy was used, and mouse identification was assigned at the time of dosing. No treatment blinding was used throughout experiments to minimise any cross-contamination between treatment and control samples, however all samples were processed and analysed in parallel. Livers were crushed in liquid nitrogen to ensure homogenous representative sample was assayed in downstream extractions. Liver, spleen, epididymal adipose and lymph nodes (pooled iliac, inguinal, axillary and mandibular) were stored at –80°C until further processing.

### RNA extraction and purification

Tissues were homogenised in QIAzol (1ml/100mg tissue) using TissueLyser LT with stainless steel beads (Qiagen) at 50Hz until large particulates were no longer visible. Homogenates and whole blood samples were centrifuged at 12,000g for 5mins to remove any insoluble material. Total RNA was isolated from QIAzol using phenol-chloroform partitioning performed according to the manufacturer protocol (Qiagen). For tissues with low RNA yield (adipose and blood), glycogen (Thermo Fisher, Cat#R0551) was used to aid in the precipitation of RNA and performed according to manufacturer protocol. An additional DNase digestion step was performed following RNA precipitation and conducted using manufacturer protocols (Merck Life Sciences, Cat# 4716728001 or NEB, Cat# M0303L). RNA was reprecipitated with the addition of 0.1x volume 3M sodium acetate, pH 5.5, followed by 1:1 addition of isopropyl alcohol and precipitated overnight at -20°C. RNA pellet was resolubilised in nuclease-free water, concentration and purity were determined using NanoDrop. RNA integrity was determined using Bioanalyzer RNA 6000 nano kit (Agilent Technologies) with samples >RIN 8 included for downstream processing. Aliquots of RNA were stored at -80°C and underwent a single freeze-thaw process. All reagents were molecular biology grade and nuclease-free where possible.

### Nanopore sequencing library preparation

Purified RNA was enriched for mRNA using NEBNext® High Input Poly(A) mRNA Isolation Module (New England Biolabs). Reverse transcription and double-stranded cDNA synthesis was performed as per the Oxford Nanopore protocol for Ligation sequencing V14 - Direct cDNA sequencing (SQK-LSK114) using reagents as per manufacturer recommendation unless otherwise specified. This single VN primer was replaced with a poly(A) binding primer pair; /5Phos/GAAGATAGAGCGACAGGCAAGT (final concentration of 2.8 µM in 10 mM Tris-HCl pH 7.5, 50 mM NaCl) and /5Phos/ACTTGCCTGTCGCTCTATCTTCTTTTTTTTTTTTTTTTTTTT (final concentration of 1.4 µM), annealed by heating to 95°C for 2 min and cooling at 0.1°C/sec. All primers were HPLC purified obtained from Integrated DNA Technologies. Mag-Bind® TotalPure NGS magnetic beads (Omega Bio-tek) were used in all protocols for sample clean-up protocols. cDNA was barcoded and sequencing libraries prepared using Oxford Nanopore Ligation sequencing amplicons - Native Barcoding Kit 96 V14 (SQK-NBD114.96) kit and protocols. Final libraries were sequenced on a PromethION 2 (P2) Solo.

### Nanopore Sequencing Analysis

The raw .pod5 files were basecalled and aligned to the GRCm39 reference genome using dorado v0.7.3 with a high accuracy basecalling model (dna_r10.4.1_e8.2_400bps_hac @v5.0.0, Oxford Nanopore Technologies). The reference genome was annotated with two additional sequences (Synth_Cre and tdTomato) to enable identification of the synthetic mRNA and induction of tdTomato expression.

A custom python script (available on GitHub) was then used to generate a read count table and analyse the degradation pattern of the synthetic mRNA. In brief, the script analyses samples on a per organ basis by extracting the sample details contained in the name of the .bam file. The naming convention of the .bam file was: treatment_biologicalreplicate_time_technicalreplicate_barcode.bam. The read count table was generated by extracting the ensemble transcript ID for each primary read in the .bam file and combining the reads from each technical replicate. The read count table was saved as a .csv for subsequent differential gene expression analysis.

### Synthetic mRNA Analysis

To determine the relative amount of synthetic mRNA in each sample, the total number of Synth_Cre reads were ratioed to the total number of endogenous reads and multiplied by 1,000,000 to determine the counts per million (cpm). To enable quantification of the mass of synthetic mRNA per gram of tissue, the cpm per ng of mRNA was determined for each tissue. A mass normalization factor was generated by spiking a known mass of synthetic mRNA (250ng for liver, spleen and lymph nodes, 50ng for adipose tissue and blood) into 100µg of tissue (liver, spleen, lymph nodes or adipose) or 200µl of blood isolated from undosed Ai14 mice. The cpm of spiked mRNA was divided by the mass of spiked mRNA per g of tissue to give the cpm per ng of mRNA per gram of tissue (summarised in Supplementary Table 8). To convert cpm to a quantitative mass of synthetic mRNA per gram of tissue, the cpm of Synth_Cre was divided by the mass normalisation factor. The reference mass normalisation factors for each tissue apply universally for quantifying the mass of any mRNA sequence present. The mass normalisation factor for each tissue should be conserved for any C57BL/6 mice, and therefore do not need to be routinely redetermined. We applied a similar approach, to determine the mass of tdTomato transcript within tissues, again using the mass normalisation factor. Adjustments were made for the higher molecular weight of the tdTomato, which we estimated based on sequence analysis to be 2,018 nucleotides.

To determine the amount of full-length synthetic mRNA and degradation profiles, the starting position of the Synth_Cre alignment was extracted from the .bam file to determine the length of the mRNA sequence. To ensure any truncation was due to mRNA degradation and not incomplete reverse transcription (or premature termination of the read by the nanopore), the 5’ sequence of the raw reads prior to the Synth_Cre alignment were analysed. Any read that was fully reverse transcribed and fully read by the nanopore is expected to have the strand switching primer sequence immediately before the Synth_Cre sequence. Any read that had <75% alignment to the strand switching primer was eliminated from the degradation analysis. The percent of full-length synthetic mRNA was determined by ratioing the number of full-length reads (which contained the strand switching primer) to the total number of Synth_Cre reads (with the strand switching primer). The calculation assumes that the efficiency of reverse transcription was the same for degraded and full-length mRNA.

To determine the degradation profile of the mRNA we generated a frequency distribution of the length of synthetic mRNA. As the mRNA sequence is reverse transcribed from the 3’ end, the nanopore sequencing will only identify the first point of degradation from the 3’ end. To take into account that a strand could be degraded in multiple positions, we applied an algorithm to calculate the probably of degradation at that base in the synthetic mRNA sequence. Where *i* is the position in the mRNA sequence, DegProb*_i_* is the probably of degradation at the i-th position in the mRNA sequence, *f_i_* is the number sequences of length *i*, and is the cumulative sum of counts from position *i* to the end of the sequence (*n*).

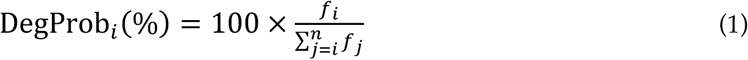

### Barcode cross-contamination

PCR-free multiplexing by ligation of unique DNA barcodes to each cDNA sample provides an efficient strategy for nanopore sequencing sample preparation (Fig 1a). While increasing throughput and reducing the sequencing cost, there is potential for barcode cross-contamination. Barcode cross-contamination can occur due to incorrectly assigned reads, or cross-reaction of barcodes during the barcoding step.

To determine the rate that barcodes are incorrectly assigned, we sequenced cDNA libraries that contained two barcoded samples, and determined the total number of reads correctly and incorrectly assigned to the 96 possible barcodes of the SQK-NBD114.96 barcoding kit. We found that barcode misassignment due to sequencing errors was rare (<1 in 10,000 reads; Supplementary Table 5).

To quantify the cross-reaction of barcodes, we performed the following analysis. In a library of multiplexed samples, the number of observed (or measured) reads for gene within a barcoded sample will be equal to the actual (or true) number of reads, minus the number of reads for that gene that have cross contaminated other barcodes, plus the number of reads for that gene that cross contaminated other barcodes. For barcode 1, this can be expressed as the equation below:

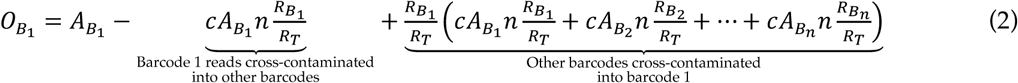

Where, 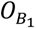 is the observed reads of a gene with barcode 1, 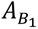 is the actual reads of the gene with barcode 1, *c* is the cross contamination rate between barcodes, *n* is the number of barcodes in the library, 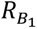 is the total number reads for barcode 1 in the library, *R_T_* is the total number reads in the library, 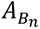 is the actual reads of the gene with barcode *n*, 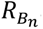 and is the total number reads for barcode 1 in the library.

To determine the cross-contamination rate (*c*), we sequenced libraries that contained only two barcodes, one sample of spleen tissue, and a second sample of spleen tissue spiked with synthetic mRNA. If there are only 2 barcodes in the library, then the observed number of reads for barcode 1 (spleen tissue only, with no synthetic mRNA) will be:

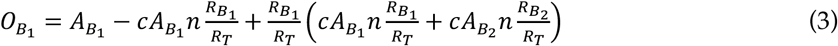

Because we know that barcode 1 contains no synthetic mRNA, 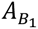 is equal to 0, so the equation simplifies to:

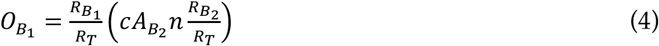

Rearranging to solve for *c* gives:

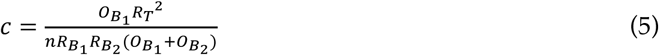

We prepared and sequenced three separate libraries with spiked and non-spiked tissue and calculated the cross-contamination rate (*c*) for each library (Supplementary Table 6).

We then used the mean cross contamination rate (0.023688) and coefficient of variance (0.16192) for subsequent corrections of the barcode cross contamination.

Equation 2 above can be written in the form of a compensation matrix, to allow unmixing of cross contamination from multiple barcodes in the one sample.

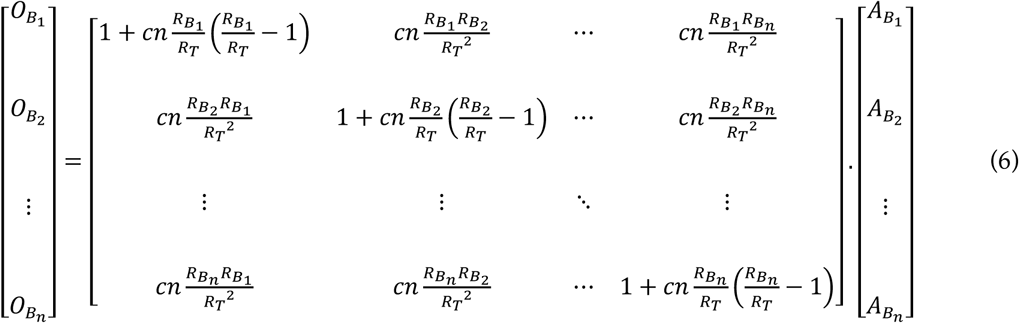

Rearranging to solve for 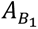 to 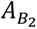 gives:

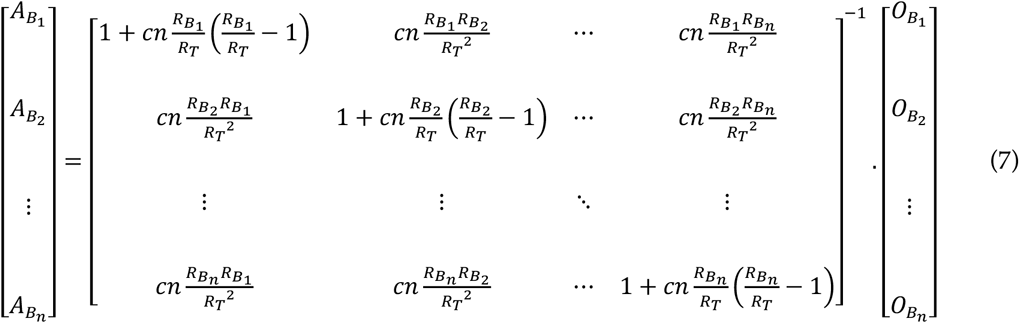

Where all terms on the right-hand side of the equation are known.

We therefore applied a compensation matrix to estimate and correct barcode cross-contamination between samples. Because this correction adds uncertainty to synthetic mRNA read counts, the LOD for compensated samples was defined as the read-depth-based LOD plus three times the error associated with the contamination correction. Applying this correction reduced false-positive Cre mRNA reads in PBS/mRNA multiplexed samples to below the LOD (Supplementary Table 7). All subsequent tissue analyses therefore incorporated barcode cross-contamination compensation.

### Differential Gene Expression Analysis

The read count table generated by the python script was loaded into R and analysed using EdgeR. The ensemble transcript ID’s were mapped to the corresponding entrez gene ID using biomaRT. All duplicate gene IDs were summed together and any transcript ID’s that did not map to a gene ID were excluded from further analysis.

Time points 1, 3, 6, 12 and 24 hours were analysed for both the LNP and PBS treated mice, along with the untreated organs as a baseline control. The samples were grouped based on the replicates (n=5 or 6). Lowly expressed genes (<15 reads across all samples) were excluded from further analysis. EdgeR’s default TMM method was used to normalise each sample.

The log2 fold change (LFC) for each gene was calculated for each time point by performing a quasi-likelihood negative binomial generalized log-linear model test (glmQLFTest), contrasting the LNP treated samples (minus the untreated control) to the PBS treated samples (minus the untreated control). Genes with a false discovery rate (FDR) of <0.05 were treated as significant.

We also performed a time course analysis of both the LNP and PBS treated samples (using the untreated control as the 0 time point for both treatment groups) by applying a cubic spline model, with a reformed spline function where first term correlates linearly with time (Z1 - indicating the overall trend in gene expression), and the second term correlates with logTime (Z2 - indicating the initial trend in gene expression). Again, genes with a false discovery rate (FDR) of <0.05 were treated as significant, and genes that were found to be significant for both PBS and LNP treatment were excluded from further analysis.

The gage package was then used to determine gene enrichment in signalling and metabolic pathways using the sigmet.idx.mm dataset. Genes not detected in the sequencing experiment were removed from each pathway to limit bias, and only pathways with >5 genes were included for analysis. We analysed genes that showed a significant change with Z1 (overall trend in gene expression) and Z2 (initial trend in gene expression). Any pathway with a p value <0.05 in either the Z1 or Z2 analysis were treated as significant.

Finally, to validate the pathways identified with the time course trend analysis, the LFC for the individual time point comparisons of LNP treatment vs PBS were extracted for each gene in the significant pathways. This data was used to generate a time course heatmap for each pathway, and a KEGG pathway map with overlayed LFC.

### RT-qPCR

We designed and evaluated several primer pairs for short amplicon targets within the Cre mRNA full sequence RT-qPCR using both Primer3web v4.1.0 (https://primer3.ut.ee/) and selected the following pair for all quantification studies; forward 5’-GTTACACAAGGGAAGAAAAGCCGC -3’, reverse 5’-CAGGTTGCTGTTCAGGTCGGTCC -3’. *Hprt1* was used as housekeeper gene which remained stable over LNP treatments; forward 5’-AGTTCTTTGCTGACCTGCTG -3’, reverse 5’-CCACCAATAACTTTTATGTCCCC -3’. To amplify the near complete Cre mRNA sequence we designed a primers pair close to each terminus which provided coverage of ∼98% of the full sequence excluding the polyA tail; forward 5’-TAATCTTGTCTCGCTCCGGG -3’ and reverse 5’-GTACGGGTGCAGGAAGGG -3’.

RNA (125ng mRNA or 1500ng extracted RNA) was reverse transcribed using AffinityScript QPCR cDNA Synthesis Kit (Agilent Technologies) following manufacturer’s recommended procedures. qPCR was performed on an AriaMx Real-time PCR System (Agilent Technologies). A 10 µl reaction volume was prepared containing 5µl SsoAdvanced™ Universal SYBR® Green Supermix (Bio-Rad, Cat# 1725272), 400 nM of each primer and 7.5ng template. For Cre mRNA cDNA serial 1:10 dilutions were made and 2.5µl template was added to each well to establish sensitivity (LOD) defined as the estimated concentration giving a Ct value lower than the mean – (2*SD) of background signal (n=3 samples). PCR efficiency (*E*) was calculated based on concentration vs. Ct slope = (10^-1/slope^ -1) x100%. For short amplicon product, cycling conditions included 95°C for 3min (initial denaturation), 40 cycles at 95°C for 5 sec and 65°C for 10 sec (annealing/extension). For full length amplicon product, cycling conditions included 95°C for 3min followed by 40 cycles at 95°C for 5 sec and 60°C for 60 sec. Melt curve was performed from 65°C to 95°C, with 0.5°C resolution, to assess product specificity. Samples were run in triplicate and raw Ct and T_m_ values were exported from Aria Mx v2.1.1 (Agilent Technologies). For tissue samples, data were expressed in Ct values normalised to *Hprt1* housekeeping gene and fold change between spiked tissue samples and unknown samples was determined using the 2^−ΔΔCt^ method and back calculated to give ng/g Cre mRNA concentration in tissue. To validate tissue detection range and limit of quantitation (LOQ) blank tissue (spleen and liver) was spiked with Cre mRNA (ng mRNA/g tissue) over a sequential 1:10 dilution series prior to RNA extraction and reverse transcription, generating a 9-point sample standard curve. qPCR was performed on three independently prepared tissue standard curves and the LOQ was determined based on the lowest Ct value which could be interpolated within >70% accuracy of the standard curve (Supplementary Table 2).

### Protein extraction, Cre recombinase and tdTomato ELISA

Tissues were sonicated (Qsonica Q125) in Radioimmunoprecipitation Assay lysis buffer (Merck, Cat# 20-188) containing cOmplete™, EDTA-free Protease Inhibitor Cocktail (Merck, Cat# 11873580001) and 25U/ml benzonase nuclease (Merck, Cat# E1014). Lysates were centrifuged at maximum speed for 15 min and supernatant collected for ELISA. Blank tissue samples from C57BL/6J mice were weighed and spiked with known amount of Cre recombinase (New England Biolabs; Cat# M0298) and tdTomato protein (Origene; Cat#TP790045) prior to sonication and were used to provide standard curves. For Cre recombinase ELISA, Nunc-Immuno™ MicroWell™ plates were coated with 2ug/ml rabbit anti-Cre antibody (Thermo Fisher; Cat#PA5-32244) in carbonate buffer (Merck; Cat #C3041-50CAP) overnight at 4°C. A blocking solution containing 5% skim-milk powder in 1x Tris buffered saline with 0.1% tween 20 (TBST) was incubated for 1h at room temperature followed by an additional block with 10% normal donkey serum (Merck; Cat#D9663) in PBS. Tissue lysates were diluted in the blocking solution and incubated overnight at 4°C. A mouse anti-Cre detection antibody (Merck; Cat#MAB3120) at a 1:1000 dilution was applied for 2h at room temperature followed by 1:20000 dilution of biotin-conjugated donkey anti-mouse IgG secondary antibody (Jackson ImmunoResearch; Cat#715-065-151) for 1h. Streptavidin-HRP at 1:1000 dilution (Jackson ImmunoResearch; Cat#016-030-084) was incubated for 30min at room temperature. A diluent of milk block was used for all steps with washing performed in triplicate with 1x TBST, as above. 1-Step™ TMB ELISA Substrate Solution (Thermo Fisher Cat#34029) was added for 10 min to allow colour development with reaction stopped with the addition of an equal volume of 2M phosphoric acid. For tdTomato ELISA a similar procedure was followed as for Cre recombinase ELISA, using 2ug/ml of goat anti-tdTomato antibody to coat plates (Origene Cat#AB8181), rabbit monoclonal anti-RFP antibody (Thermo Fisher; Cat# 600-401-379) was used at 1:5000 for detection, and biotin-conjugated donkey anti-rabbit IgG (Jackson ImmunoResearch; Cat# 711-065-152) was used as secondary antibody at 1:20000 dilution.

Absorbance was read at 450nm on a BMG CLARIOstar plate reader and samples concentrations were interpolated from sigmoidal 4-parameter logistic nonlinear regression curves using GraphPad Prism 10. The LOD was defined as the protein concentration in tissue giving an absorbance value greater than the mean + 3*(SD) of background tissue signal (n=3 samples). For liver and spleen, triplicate standard curves were prepared on different days with lower limit of quantitation (LLOQ) and the upper limit of quantification (ULOQ) defined as the lowest and highest standard, respectively, with a %backfit of 75%-125% and %CV ≤ 20% (Supplementary material Table 9).

### Cytokine multiplex assay

An additional cohort of Ai14 mice were dosed intravenously with 0.5 mg/kg Cre mRNA encapsulated LNP or PBS and randomised across time points for spleen and plasma collection to assay cytokines and chemokines. Plasma was isolated within 10mins of blood collection by centrifugation at 1000xg for 10mins and snap frozen on dry ice. Tissue was homogenised and prepared according to Thermo Fisher lysis protocol for Luminex assays. Briefly, spleens were homogenised in 0.5ml/100mg in Cell Lysis Buffer (Cat. No. EPX-99999-000) using the Tissue Lyser LT at 25 Hz. Following centrifugation at 16,000 × g for 10 mins at 4°C, supernatants were removed and protein concentration determined by BCA assay (Pierce™ BCA Protein Assay Kit, Thermo Fisher; Cat# 23227). Samples were diluted to 10mg/ml protein in PBS and processed for ProcartaPlex™ Mouse Cytokine & Chemokine Convenience Panel 1A, 36plex (Thermo Fisher; Cat# EPX-88182-000) following manufacturer guidelines. The samples were run in duplicate on a Luminex MAGPIX® and data analysed using ProcartaPlex Analysis App (Thermo Fisher) to give cytokine and chemokine concentrations. Where a concentration could not be determined (typically in PBS dosed control samples) as it was below the assay quantification, the lower limit of quantitation for the assay was used as a substitute value to enable fold change calculation to be determined.

### Immunohistochemistry

Spleen and liver tissue was fixed overnight in 4% paraformaldehyde in PBS before cryoprotection in a sucrose gradient up to 30% in PBS, prior to snap freezing in isopentane over dry ice. Tissue was cryosectioned at 20µm thickness and air-dried onto poly-L-coated glass slides. Sections were blocked in 6% normal donkey serum (Jackson ImmunoResearch; Cat# 017-000-121) with 0.3% (v/v) Triton-X 100 in PBS for 1h at room temperature. The following primary antibodies were prepared in 1% serum: 0.3% triton-X in PBS; 1:200 rat anti-F4/80 (Abcam; Cat# ab6640); 1:500 rabbit anti-ASGR1 (SinoBiological; Cat# 50083-R114) and 1:100 hamster anti-CD11c (ThermoFisher; Cat# 14-0114-82) and incubated with tissue sections overnight at 4°C. Following washing 3x in PBS, sections were incubated for 2h at room temperature in the following secondary antibodies prepared at 1:400 dilution; donkey anti-rabbit Alexa Fluor®-488 (Jackson ImmunoResearch; Cat# 711-545-152); donkey anti-rat Alexa Fluor®-647 (Jackson ImmunoResearch; Cat# 712-605-153); donkey anti-hamster Alexa Fluor®-488 (Jackson ImmunoResearch, Cat# 127-545-160). All images were captured on a Leica SP8 Confocal.

### Flow cytometry

At 24 hours post LNP administration, blood and tissues were collected and dissociated into single cell suspension for immunostaining as previously described^36^. All immune cell pellets were stained with a flow cytometry panel containing the following antibodies: αCD3e-BV650 mAb (clone 145-2C11, BD Biosciences), αCD19-BV786 mAb (clone 1D3, BD Biosciences), αCD11b-BV421 mAb (clone M1/70, BioLegend), αLy-6C-BUV661 mAb (clone HK1.4.rMAb, BD Biosciences), αCD49b-FITC mAb (clone DX5, BD Biosciences), αLy-6G-BV605 mAb (clone 1A8, BD Biosciences), αCD45-Pacific Blue mAb (clone S18009F, BioLegend), αI-A/I-E-BV510 mAb (clone M5/114.15.2, BioLegend), αF4/80-PE/Dazzle mAb (clone BM8, BioLegend), and αCD11c-Alexa Fluor 700 mAb (clone N418, BioLegend). Additionally, Mouse BD Fc Block™ and viability dye (LIVE/DEAD™ Fixable Blue Dead Cell Stain Kit, Thermo Fisher) were included. Samples were incubated on ice for 30 minutes, followed by washing to remove excess antibody.

Flow cytometry was performed using a Cytek Aurora 5 laser cytometer, and data were analyzed using FlowJo (BD Biosciences). Leukocyte phenotyping was conducted using the following markers: CD3+ T cells (CD45+, CD11b-, CD3e+,); dendritic cells (CD45+, CD3e-, CD19-, CD11c+, MHCII+); monocytes (CD45+, CD11b+, Ly6C+, Ly6G-); neutrophils (CD45+ CD11b+, Ly6C+, Ly6G+); macrophages (CD45+ CD11b+, Ly6C low, Ly6G-, F4/80+, SSA low); and CD19+ B cells (CD45+, CD11b-, CD3-, CD19+ or MHCII+) with gating strategy illustrated for liver, spleen, lymph node and blood in Supplementary Figure 4. Percentage cell populations are reported as mean ± SD, n=3 animals.

### Statistical Analysis

Statistical analysis was performed as detailed below using GraphPad Prism 10.0 with outcomes provided within each figure legend. Cre mRNA integrity assessments comparing nanopore sequencing with microfluidic electrophoresis compared mRNA pre and post formulation into LNPs with two-sided paired t-test, with pairing between each mRNA and LNP encapsulated replicate. Area under curve (AUC) was calculated using GraphPad prism which generated a standard error (SE). The degrees of freedom (df) within each tissue were defined as the number of data points for the tissue subtracted by the number of time points. The n was defined as df + 1. AUC for Cre mRNA biodistribution over time was compared using Welch one-way ANOVA (due to significantly different SD between tissues). Following a significant ANOVA (P<0.05), Dunnett’s post comparison was performed comparing all tissues to liver, the primary organ of LNP uptake. AUCs for Cre protein, tdTomato mRNA and protein biodistributions were also calculated using GraphPad prism. Prior to AUC calculation a significant elevation in tdTomato expression above baseline was confirmed: for transcript this required P<0.05 for two-way ANOVA interaction effect between treatment and time, for protein P<0.05 for one-way ANOVA comparing time points with t0. The average baseline mRNA and protein level of tdTomato were then subtracted from all respective time points and AUC calculated using GraphPad prism. The ratio of average AUCs (R) of protein (A) to mRNA (B) was calculated (R=A/B) for Cre and tdTomato. To calculate the absolute error of R (SE_R_), error of propagation formula was used, whereby the relative error (RE_R_) was calculated by taking the square root of the sum of relative errors where the relative error is the SE divided by the AUC value; √ [(SE_A_/A)^2^ + (SE_B_/B)^2^]. SE_R_ is then obtained by multiplying the RE_R_ by the ratio R.

For cytokine protein concentrations measured in spleen, one-way ANOVA was performed to indicate a significant change over time. Fold change was determined against the average cytokine concentration measured in PBS control group, a log_2_ transformation was applied to all data values before averaging each time point. The strength of Log_2_-fold changes (LFC) in transcript correlating to protein were compared by Pearson’s correlation coefficient (r) generated by Prism, where we considered r>0.6 a strong correlation and r=>0.8 a very strong correlation.

## Data availability

Source data can be provided with this paper. The transcriptomic datasets generated in the current study will be made publicly available in the Zenodo repository, upon publication of this article. Requests for additional materials should be addressed to A.P.R.J.

## Code availability

The code used to generate all analyses and figures in this study is available in our GitHub repository (https://github.com/AngusNano/Nanopore_biodistribution/tree/main).

## Acknowledgements

A.P.R.J. was supported by an NHMRC Career Development Fellowship (GNT1141551) as well as ARC Discovery Projects (DP210103174) and NHMRC Ideas Grant (GNT2011963). This research was also partially funded by the Victoria State Government through funding support from mRNA Victoria for the Victorian mRNA Innovation Hub.

## Funding

Open access funding provided by Monash University

## Contributions

A.P.R.J. conceptualized the study and designed the research framework. A.P.R.J. and V.M.M. developed the methodology. V.M.M., M.Z.C., B.R.H., Y.Y., L.M., P.K., T.J.P., S.A.F. conducted experimental investigations. A.P.R.J., V.M.M., O.M.F., and C.J.H.P. performed data analysis. A.P.R.J., V.M.M., D.Y., and S.A.B. conducted data visualization and figure design. A.P.R.J., V.M.M., O.M.F., and C.J.H.P. conducted data analysis. The project was supervised by A.P.R.J., C.W.P., C.J.H.P. The original draft was written by A.P.R.J., V.M.M., and D.Y., with critical revisions and editing contributed by C.J.H.P. All authors reviewed and approved the final manuscript.

## Ethics declarations

### Competing interests

The authors declare no competing interests.

## Notes

### Competing Interest Statement

The authors have declared no competing interest.

