## Supplementary Fig. 1a-h for "Long-read sequencing quantifies synthetic mRNA abundance, integrity and host response in vivo"

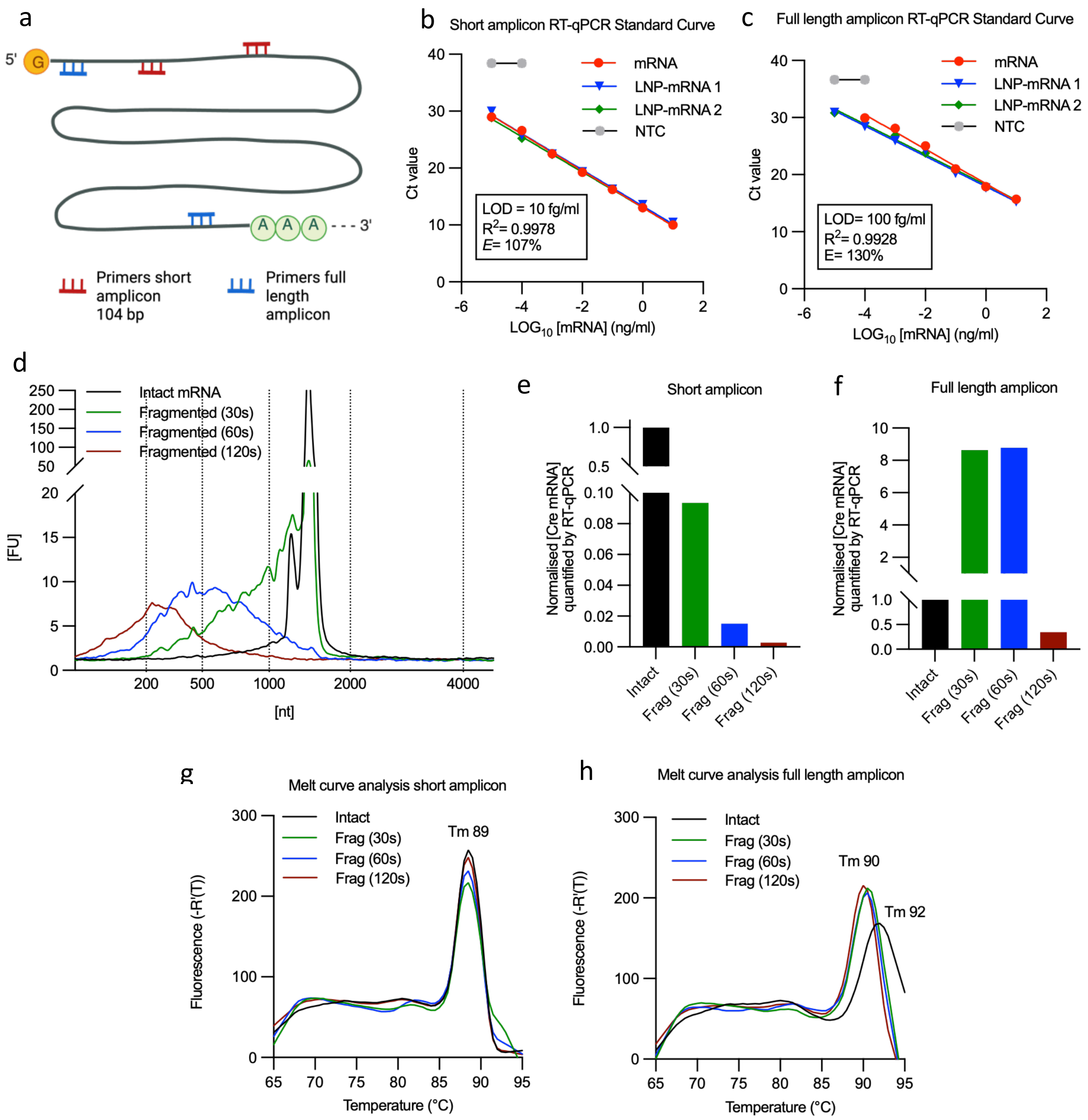


**Supplementary Figure 1. Design and validation of RT-qPCR primers against Cre recombinase mRNA. a**, RT-qPCR primers were designed to amplify a short 104 base pair region within the 5′ UTR and CDS region of Cre recombinase, and a much longer 1271 base pair amplicon representing majority of the Cre recombinase sequence from 5′ to 3′-UTRs. **b**, Cre recombinase IVT mRNA was reverse transcribed (triplicate samples were prepared with 2 being released from LNP encapsulation), and a dilution series was amplified by RT-qPCR using short amplicon conditions to generate a standard curve of cycle threshold (Ct) vs. mRNA concentration. **c**, Samples were also amplified using full-length amplicon conditions. A non-template control (NTC) was included which did not contain the cDNA template. The limit of detection (LOD), $R^2$ of standard curve and amplification efficiency ($E$) are displayed. **d**, Electropherogram output showing the product size for Cre mRNA following simulated fragmentation with divalent cation-mediated hydrolysis with almost complete abolishment of full-length product following ≥60s incubation. Quantification of Cre mRNA present in samples generated in d based on RT-qPCR **e,** short amplicon and **f,** full-length amplicon conditions, using standard curve interpolation and normalisation to the intact unfragmented mRNA. The melt curve analysis was performed for the amplicons generated for both amplification conditions (e, f), **g,** showing good alignment for melting temperature for all products generated using short amplicon conditions regardless of the degree of fragmentation, **h,** in contrast the reduction in melting temperature shows a different product is being amplified in the fragmented samples when using the full-length amplicon conditions.

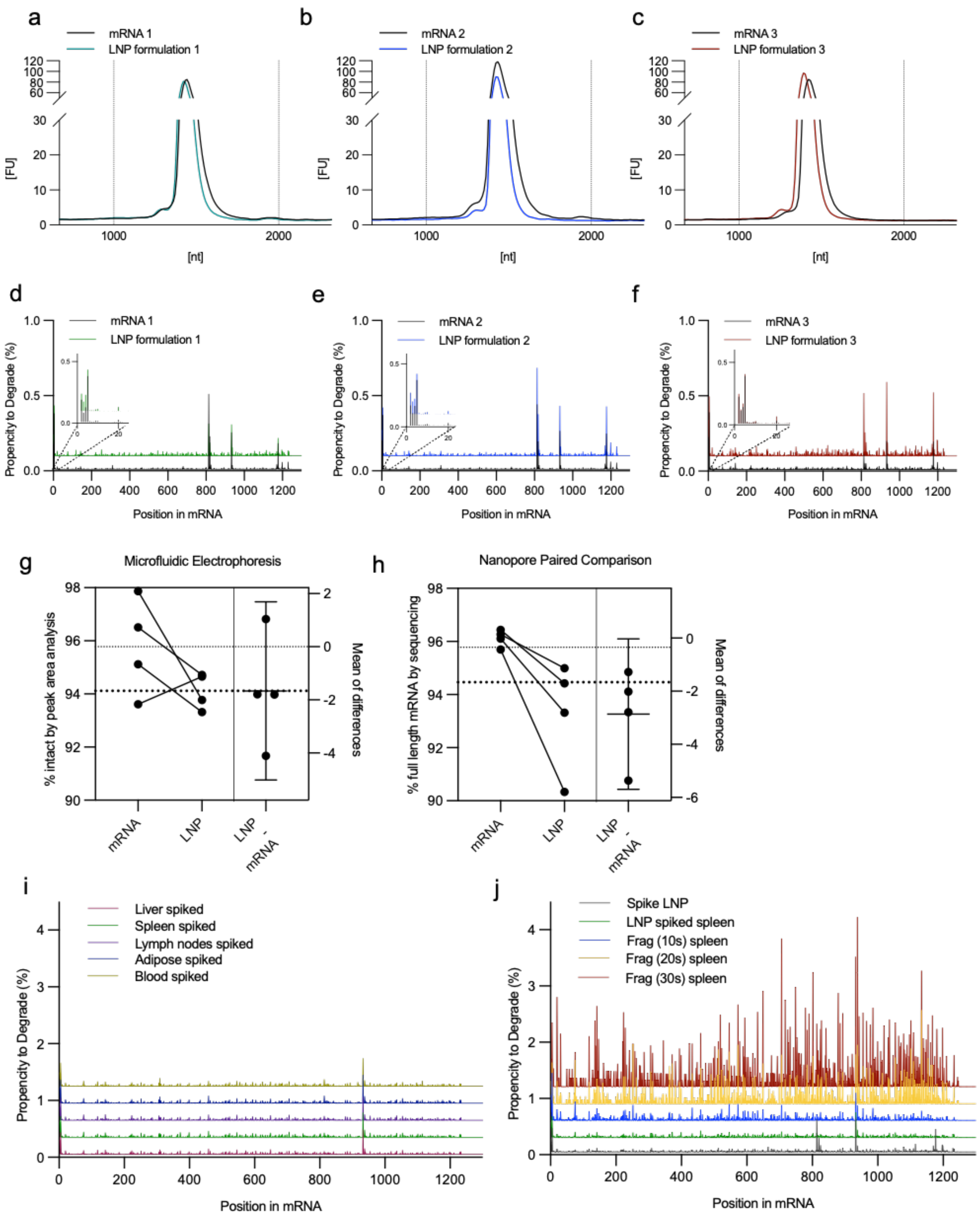


**Supplementary Figure 2. Cre mRNA integrity following encapsulation into lipid nanoparticles and processing of tissue for RNA extraction**. **a-c**, Electropherogram output showing fluorescence intensity against mean product size for Cre mRNA following LNP encapsulation in triplicate formulations prepared for administration. **d-f**, Positional representation of the identical samples (a-c) across the full Cre mRNA sequence from 5' to 3' with the % propensity to a show a breakage at that nucleotide location. Inset shows there is higher propensity for breakage within the initial nucleotides of the 5' UTR. **g**, Pairwise analysis of the primary peak area as a percentage of total area in electropherogram trace for data generated in a-c and main text fig 1b comparing mRNA pre and post LNP encapsulation.

**h**, Pairwise analysis of the full length reads as a percentage of total reads for data generated in d-f and main text fig 1c comparing mRNA pre and post LNP encapsulation. Mean of difference data was generated by GraphPad prism. **i**, Positional representation across the full Cre mRNA sequence from 5' to 3' with the % propensity to a show a breakage at each nucleotide location following spiking of mRNA encapsulated LNP into tissues prior to RNA extraction. **j**, Positional representation of the % propensity to a show a breakage at each nucleotide location following spiking of spleen tissue with fragmented Cre mRNA prior to RNA extraction protocol. Data for LNP spiked spleen is replicated from i to provide a reference for unfragmented spiked spleen.

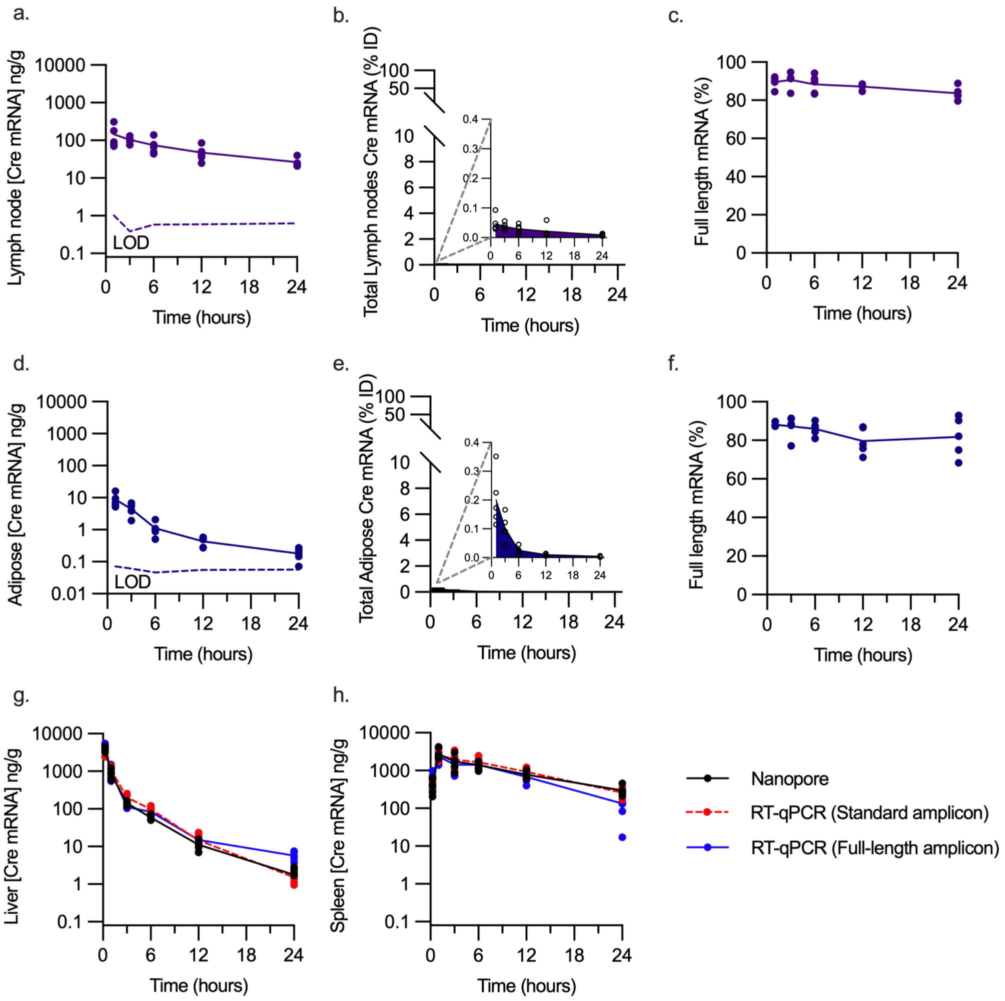


**Supplementary Figure 3**. **Time course of Cre mRNA in low yield tissues and comparison of nanopore sequencing with RT-qPCR quantification.** Biodistribution of Cre mRNA as **a**, mass per gram of lymph node, **b**, %ID within pooled lymph nodes sampled with inset showing enhanced y-axis range, and **c**, percentage of Cre mRNA represented as full length in lymph nodes (n=5 mice all time points). Biodistribution of Cre mRNA as **d**, mass per gram of adipose, **e,** %ID within total adipose (determined as 11 or 12% of total body weight in male and female mice, respectively) with inset showing enhanced y-axis range, and **f**, percentage of Cre mRNA represented as full length in lymph nodes (n=5 mice all time points). **g**, Time course biodistribution of Cre mRNA as mass per gram of liver tissue for extracted RNA samples processed for nanopore sequencing (data replicated from fig 2f) and processed for RT-qPCR using both standard amplicon and full-length amplicon conditions. **h,** Time course biodistribution of Cre mRNA as mass per gram of spleen tissue as described in g.

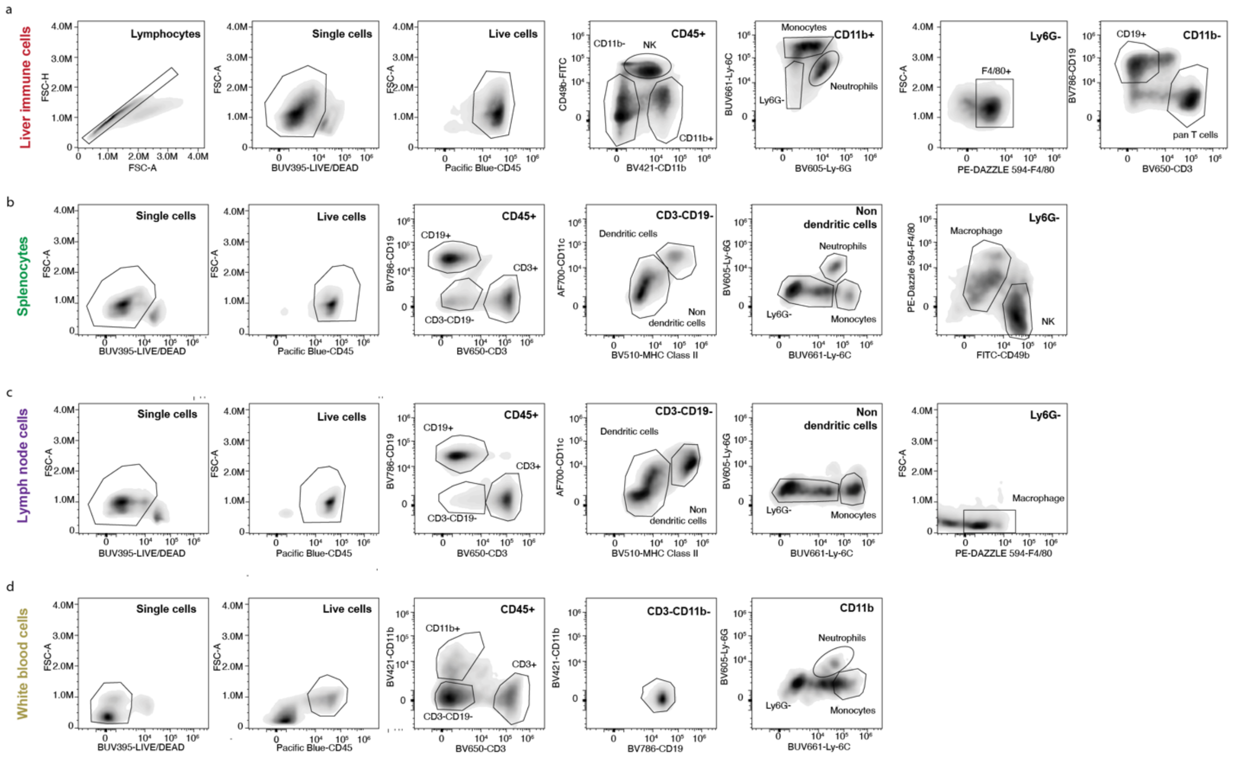


**Supplementary Figure 4. Flow cytometry gating strategy to identify subpopulations of immune cells with tdTomato expression**. **a**, Liver immune cells **b**, splenocyte suspension, **c**, lymph node suspension and **d**, white blood cells were stained with an antibody cocktail containing Live and Dead dye, anti-CD45, anti-CD49b, anti-CD11b, anti-Ly6C, anti-Ly6G, anti-F4/80, anti-CD19, anti-CD3, anti-CD11c and anti-MHCII. The cell populations were gated based on FSC-A vs SSC-A and live cells gated as the population negative for the live/dead dye. Immune cells were determined as being positive for CD45. All other subpopulations including monocytes, macrophages, neutrophils, dendritic cells (DC), T and B lymphocytes and natural killer (NK) cells were gated based on their cell surface markers.

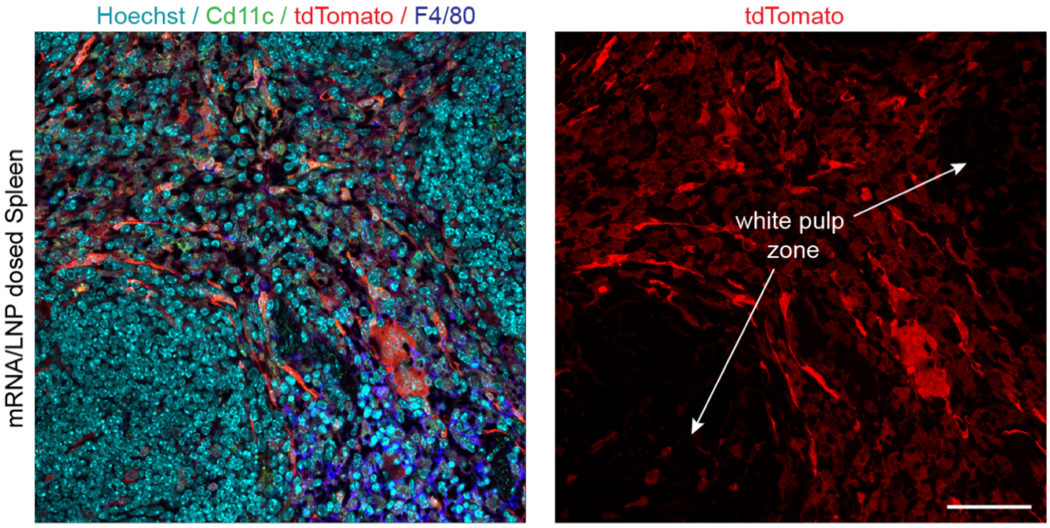


**Supplementary Figure 5. tdTomato fluorescence distribution within mouse spleen tissue section.** Overlay of tdTomato fluorescent protein expression with immunostaining for nuclei (Hoechst stain), macrophages (F4/80) and dendritic cells (CD11c) in a spleen section taken 6h post administration of Cre mRNA/LNP. Right panel shows tdTomato fluorescence only where the densely packed white pulp zones can be observed to be largely void of positive cells. Scale bar = 50µm.

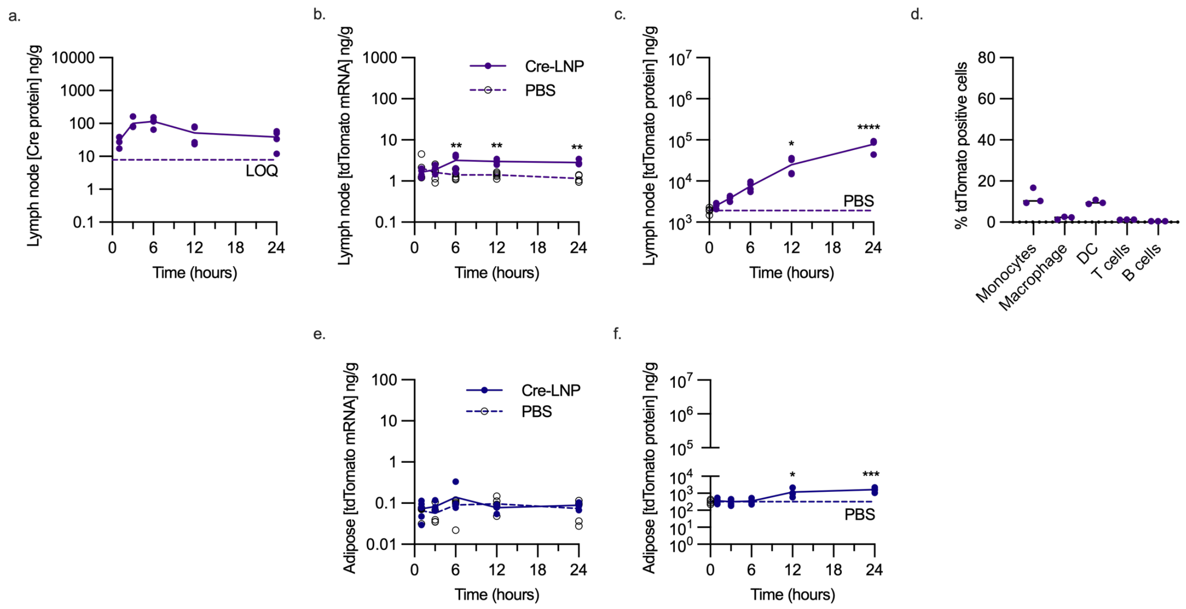


**Supplementary Figure 6. Biodistribution of Cre protein, tdTomato transcript and protein expression in LN and adipose following mRNA/LNP administration. a**, Quantification of Cre recombinase protein by ELISA presented as mass of Cre recombinase per gram of lymph node over 24h, n=4 mice per time point. The limit of quantitation (LOD) determined in undosed spiked lymph node tissue standards is displayed as dashed line (n=3). **b**, Quantification of tdTomato reporter gene transcript over time in lymph node following Cre mRNA/LNP (filled symbol, solid line) or PBS dosed mice (open symbol, dashed line); **P<0.01, by two-way ANOVA with Sidak's post comparison between LNP and PBS groups, n=5 mice per time point. **c**, Quantification of tdTomato protein expression in lymph node by ELISA following Cre mRNA/LNP with the baseline expression determined in PBS control mice (open symbol at time=0, dashed line representing mean; n=4 mice per time point). *P<0.05, ****P<0.0001 by one-way ANOVA with Dunnett's post comparison test comparing all time points to t0. **d**, tdTomato positive expression in immune cells of the liver at 24h post administration of Cre mRNA/LNP assessed by flow cytometry (n=3 mice). **e**, Quantification of tdTomato reporter gene transcript expression in adipose (solid symbol and line) and PBS (open symbol, dashed line), n=5 mice per time point. **f**, tdTomato protein expression in adipose over time (n=4 mice per time point), with baseline expression represented by PBS mice (t0, open symbol, dashed line representing mean). *P<0.05, ***P<0.001 by one-way ANOVA with Dunnett's post comparison test comparing all time points to t0.

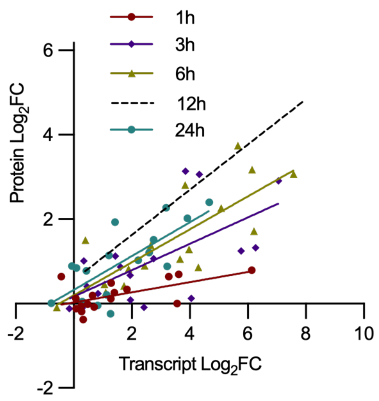


**Supplementary Figure 7. Transcript and protein expression change correlation.** Correlation between cytokine transcript and protein log2-fold changes in spleen at 1, 3, 6 and 24 h. Dotted line represents correlation at 12h which appears in main text Fig 4b. Pearson correlation *r* values were 0.6035, 0.6377, 0.7945, and 0.7363 at 1, 3, 6 and 24 h, respectively with P values of 0.0103, 0.0059, 0.0001 and 0.0008 indicating a positive correlation across all time points.

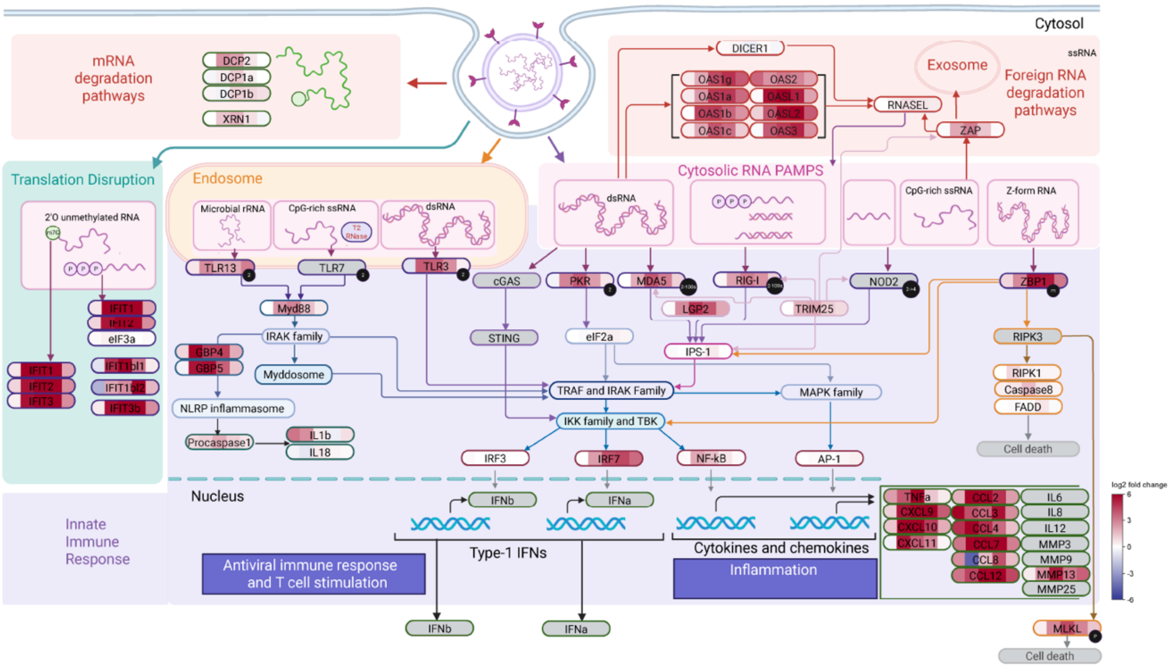


**Supplementary Figure 8. Graphical representation of signalling pathways found to be upregulated following LNP-mediated mRNA delivery to the liver.** The heatmap overlaying the gene name indicates relative upregulation at 1, 3, 6, 12 and 24h (based on Log2-fold change compared to the PBS control).

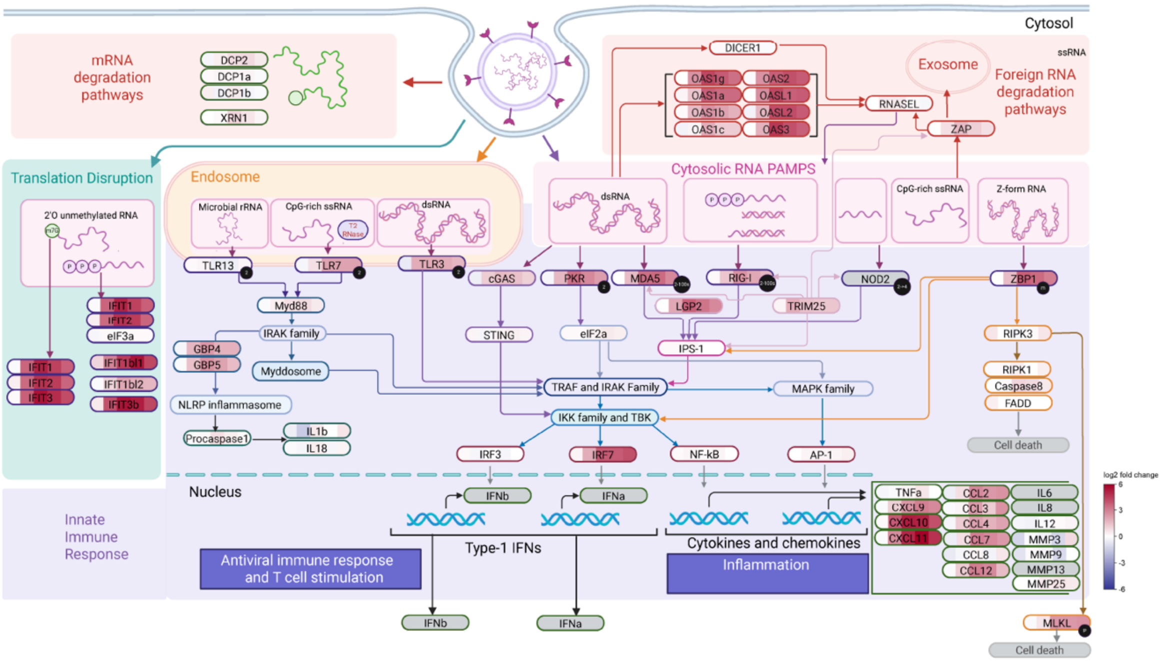


**Supplementary Figure 9. Graphical representation of signalling pathways found to be upregulated following LNP-mediated mRNA delivery lymph nodes (inguinal, iliac, axillary and mandibular)**. The heatmap overlaying the gene name indicates relative upregulation at 1, 3, 6, 12 and 24h (based on Log2-old change compared to the PBS control).

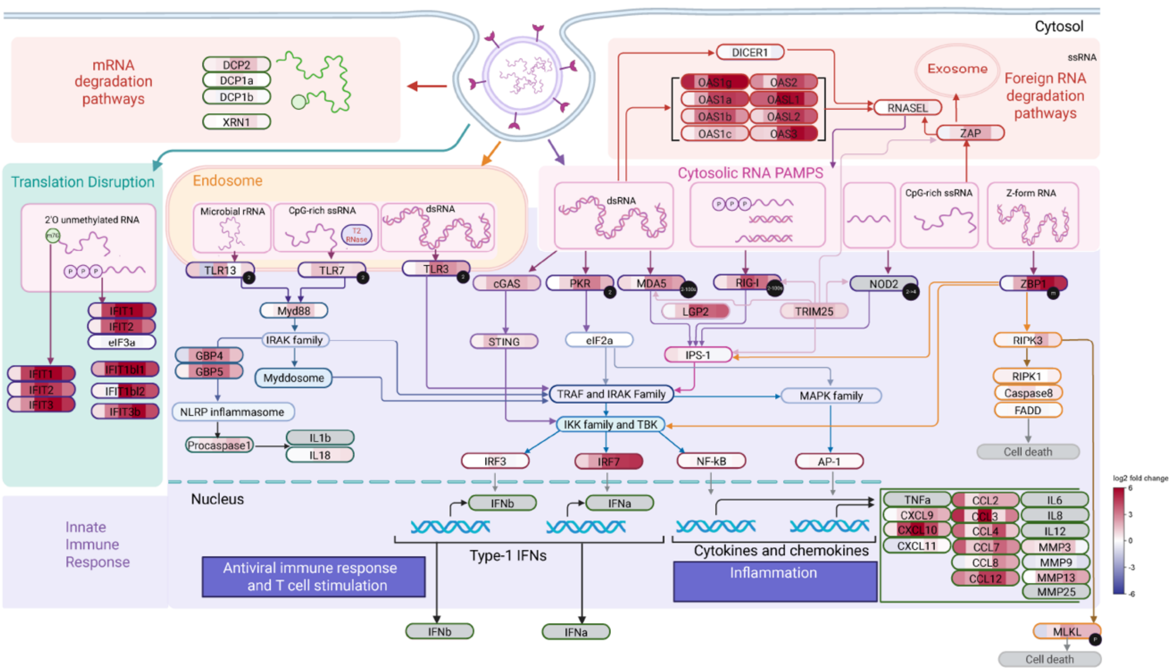


**Supplementary Figure 10. Graphical representation of signalling pathways found to be upregulated following LNP-mediated mRNA delivery to adipose tissue**. The heatmap overlaying the gene name indicates relative upregulation at 1, 3, 6, 12 and 24h (based on Log2-fold change compared to the PBS control).

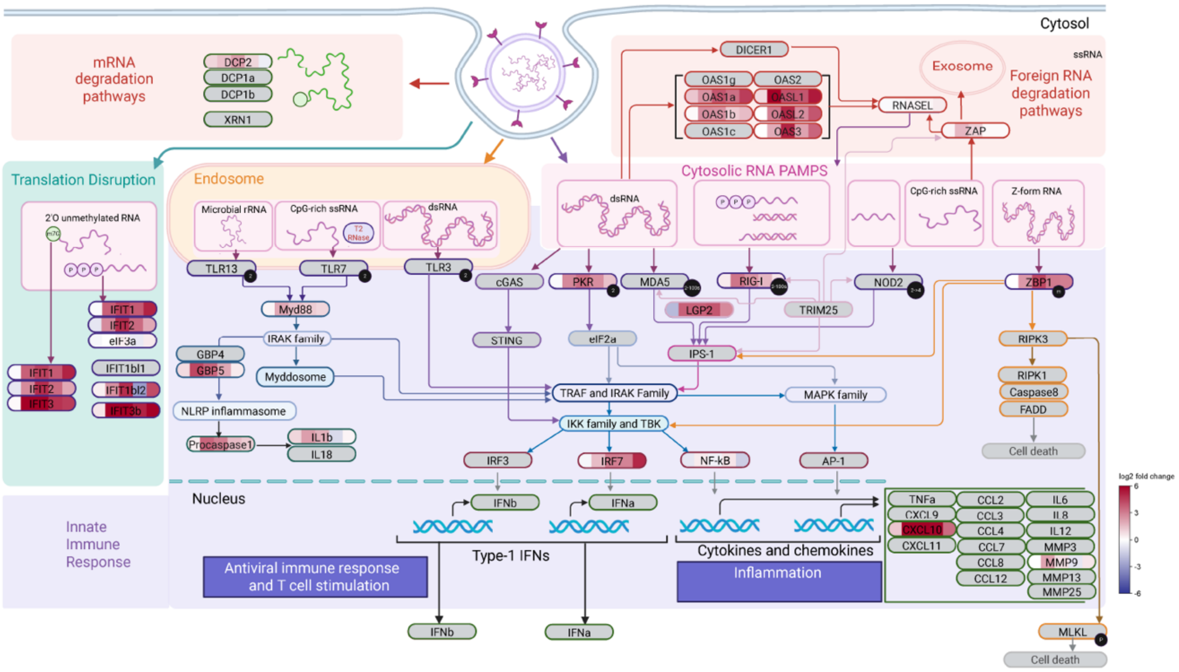


**Supplementary Figure 11. Graphical representation of signalling pathways found to be upregulated following LNP-mediated mRNA delivery to blood**. The heatmap overlaying the gene name indicates relative upregulation at 1, 3, 6, 12 and 24h (based on Log2-fold change compared to the PBS control).

**Supplementary Table 1. Determination of endogenous mRNA content of different tissues using synthetic mRNA as a reference**

| Tissue | mRNA content (ng/g) | CV (%) |
|---|---|---|
| Blood | 5.3 | 10.8 |
| Liver | 158 | 12.8 |
| Spleen | 96.7 | 9.7 |
| Lymph node | 79.6 | 11.5 |
| Adipose | 4.7 | 19.8 |

**Supplementary Table 2. RT-qPCR Assay Validation for Cre mRNA in Tissues**

| | **Liver** | | **Spleen** | |
|---|---|---|---|---|
| **Primer pair** | Short amplicon | Full length amplicon | Short amplicon | Full length amplicon |
| **Linear dynamic range** | 0.005ng/g - 5000ng/g | 0.5ng/g - 5000ng/g | 0.05ng/g - 5000ng/g | 0.5ng/g - 5000ng/g |
| **$R^2$** | 0.9972 | 0.9925 | 0.9952 | 0.9929 |
| **LLOQ** | 0.02 ng/g | 2 ng/g | 0.05 ng/g | 20 ng/g |
| **Ct at LLOQ** | 27.8 | 28.5 | 28.1 | 26.7 |
| **(%CV) of LLOQ Ct** | 1.35% | 2.36 % | 1.73 % | 5.61 % |

LLOQ, lower limit of quantitation; Ct, cycle threshold

**Supplementary Table 3. Non-linear regression analysis of Cre mRNA elimination from tissue using exponential two-phase decay modelling**

| **Organ** | **Fast phase** | | **Slow phase** | | **$R^2$** |
|---|---|---|---|---|---|
| | % | $T_{1/2}$ (min) | % | $T_{1/2}$ (min) | |
| Blood | 90.3 | 4.6 | 9.8 | 235 | 0.67 |
| Liver | 95.4 | 17.2 | 4.6 | 142 | 0.98 |
| Spleen | - | - | 100 | 303 | 0.82 |
| Lymph nodes | - | - | 100 | 260 | 0.57 |
| Adipose | - | - | 100 | 102 | 0.81 |

**Supplementary Table 4. Comparison of Long-read sequencing and RT-qPCR tissue quantification of Cre mRNA**

| | **$AUC_{0-24}$ (ng/g.h) Liver Cre mRNA** | **$AUC_{0-24}$ (ng/g.h) Spleen Cre mRNA** |
|---|---|---|
| **Nanopore** | 3237 ± 326 | 23,341 ± 2414 |
| **RT-qPCR standard** | 3959 ± 430 | 26,105 ± 3435 |
| **RT-qPCR full length** | 3936 ± 488 | 20,396 ± 2657 |

Mean ± SE calculated in Prism

**Supplementary Table 5. Incorrect barcode assignment rate**

| | Correct barcode | Incorrect barcode | Incorrect ratio |
|---|---|---|---|
| A | 3440475 | 64 | 1.86021E-05 |
| B | 2652095 | 71 | 2.67713E-05 |
| C | 2585532 | 77 | 2.97811E-05 |

**Supplementary Table 6. Barcode cross-contamination rate**

| Treatment | Library Preparation | Barcode | Total Reads | Synthetic Reads | Cross contamination (c) |
|---|---|---|---|---|---|
| Spiked with synthetic | A | barcode20 | 1747986 | 23477 | - |
| No synthetic | A | barcode25 | 1692489 | 319 | 0.02681821 |
| Spiked with synthetic | B | barcode21 | 1603928 | 22443 | - |
| No synthetic | B | barcode24 | 1048167 | 282 | 0.02595841 |
| Spiked with synthetic | C | barcode23 | 1102718 | 14735 | - |
| No synthetic | C | barcode26 | 1482814 | 133 | 0.01828596 |

**Supplementary Table 7. Barcode Contamination Correction**

| Sample Details | Total Reads | Unaligned Reads | Genomic Reads | Synthetic Reads | Barcode Corrected Synthetic Reads | LOD | Above LOD |
|---|---|---|---|---|---|---|---|
| pbs_24_A | 4858782 | 512836 | 4345830 | 116 | 4 | 69 | FALSE |
| dmg_12_A | 6840520 | 728484 | 6111424 | 612 | 469 | 90 | TRUE |
| dmg_12_B | 5294048 | 676216 | 4617368 | 464 | 351 | 71 | TRUE |
| dmg_12_C | 4295922 | 572446 | 3723214 | 262 | 166 | 59 | TRUE |
| dmg_12_D | 3292226 | 438660 | 2853210 | 356 | 285 | 45 | TRUE |
| dmg_1_B | 6391256 | 707420 | 5652998 | 30838 | 31604 | 392 | TRUE |
| dmg_1_C | 4028530 | 469116 | 3546146 | 13268 | 13420 | 86 | TRUE |
| dmg_1_D | 6190522 | 649970 | 5517144 | 23408 | 23936 | 275 | TRUE |
| dmg_24_A | 5902038 | 669982 | 5231858 | 198 | 64 | 83 | FALSE |
| dmg_24_B | 5791006 | 729516 | 5061302 | 188 | 56 | 81 | FALSE |
| dmg_24_C | 4352660 | 613896 | 3738610 | 154 | 55 | 61 | FALSE |
| dmg_24_D | 4830378 | 566828 | 4263362 | 188 | 79 | 68 | TRUE |
| dmg_3_A | 3592356 | 477442 | 3112552 | 2362 | 2317 | 33 | TRUE |
| dmg_3_B | 4915798 | 629956 | 4281898 | 3944 | 3918 | 27 | TRUE |
| dmg_3_C | 6100522 | 650570 | 5444564 | 5388 | 5397 | 23 | TRUE |
| dmg_3_D | 3854262 | 456732 | 3394538 | 2992 | 2955 | 30 | TRUE |
| dmg_6_A | 7026744 | 780642 | 6243984 | 2118 | 2021 | 69 | TRUE |
| dmg_6_B | 4785826 | 573144 | 4211194 | 1488 | 1408 | 53 | TRUE |
| dmg_6_C | 6191536 | 752794 | 5436138 | 2604 | 2533 | 53 | TRUE |
| dmg_6_D | 6233758 | 819102 | 5412680 | 1976 | 1886 | 62 | TRUE |

**Supplementary Table 8. Mass normalisation factor determined for each tissue by calibrating fixed mass amounts of Cre mRNA into tissue and quantifying the ratio of synthetic to endogenous mRNA reads**

| Tissue | Mass Normalisation Factor (cpm.ng$^{-1}$.g) | Standard Deviation | CV (%) |
|---|---|---|---|
| Blood | 219.3 | 20.9 | 9.52 |
| Liver | 9.59 | 0.98 | 10.2 |
| Spleen | 16.49 | 1.34 | 8.13 |
| Lymph Nodes | 19.93 | 2.07 | 10.4 |
| Adipose | 328.9 | 60.2 | 18.3 |

Counts per million (cpm), standard deviation based on n=6 independently spiked mouse tissue.

**Supplementary Table 9. ELISA assay Validation for Cre recombinase and tdTomato protein in tissues**

| | Liver | | Spleen | | Lymph node | | Adipose | Plasma |
|---|---|---|---|---|---|---|---|---|
| Protein | Cre | tdTomato | Cre | tdTomato | Cre | tdTomato | tdTomato | tdTomato |
| LOD (ng/g) | 0.49 ± 0.0 | 0.26 ± 0.11 | 1.63 ± 0.28 | 0.26 ± 0.11 | 5.21 ± 2.26 | 0.16 ± 0.11 | 0.059 ± 0.033 | 0.046 ± 0.030 |
| LLOQ (ng/g) | 1.95 | 0.39 | 1.95 | 0.39 | 7.81 | 0.39 | 0.16 | 0.078 |
| ULOQ (ng/g) | 125 | 25 | 125 | 12.5 | 125 | 25 | 10 | 5 |
| %CV of LLOQ | 19.4 | 1.24 | 6.46 | 14.2 | 8.50 | 7.01 | 7.37 | 9.98 |
| % accuracy at LLOQ | 92 ± 18 | 96 ± 1.2 | 111 ± 7.2 | 95 ± 14 | 111 ± 9.4 | 99 ± 7.0 | 102 ± 2.8 | 96 ± 6.5 |

LOD, limit of detection; LLOQ, lower limit of quantitation; ULOQ, upper limit of quantitation
